# A context dependent hierarchy of APOBEC3A and APOBEC3B mutators in lung adenocarcinoma

**DOI:** 10.64898/2026.03.09.710641

**Authors:** Josefine Striepen, Luka Culibrk, Alexandra Dananberg, Jacob S. Rozowsky, Mia Petljak, John Maciejowski

## Abstract

APOBEC3 cytosine deaminases are major sources of cancer mutations. In lung adenocarcinoma (LUAD), APOBEC3 activity drives tumor evolution and therapy resistance, nominating these enzymes as therapeutic targets. Although APOBEC3A drives the major portion of APOBEC3-associated mutational signature burdens across cancers, the relative contributions of individual APOBEC3 paralogs in LUAD remain unclear, limiting effective targeting. Here, we define the roles of endogenously misregulated APOBEC3 deaminases in LUAD using CRISPR–Cas9 knockouts and whole-genome sequencing of 197 long-term–propagated single-cell clones. We show that APOBEC3A and APOBEC3B both generate mutations but display marked heterogeneity, ranging from single-enzyme to dual activity or minimal mutagenesis, which is not predicted by commonly used mRNA and protein levels, or deaminase assays. Using these genetically defined systems, we provide the first causal evidence that endogenous APOBEC3A and APOBEC3B generate the indel signature InD9a in tumors, linking it to uracil excision following cytosine deamination. Together, these findings reveal heterogeneous APOBEC3 contributions to LUAD mutagenesis and highlight the need for biomarkers that resolve individual APOBEC3 activities for their effective targeting.

## Introduction

Cytosine deamination by APOBEC3 enzymes is a major source of mutations in cancer, generating the genome-wide single base substitution (SBS) mutational signatures SBS2 and SBS13 (SBS2/13) in >80% of all tumor types^1,2^. SBS2 and SBS13 are characterized by C>T and C>G/C>A mutations, respectively, at TC dinucleotides^3^. APOBEC3 mutational signatures are particularly prominent in LUAD, occurring in >50% of tumors, with multiple recent independent lines of evidence implicating APOBEC3 activity in LUAD evolution and therapy resistance^4–7^. Although APOBEC3 deamination represents a major mutational force in LUAD, the contributions of individual APOBEC3 enzymes to mutagenesis remain incompletely understood.

Substantial data implicates APOBEC3A and APOBEC3B paralogs as the principal sources of APOBEC3-associated mutations in cancer. APOBEC3B is frequently highly expressed in tumors, and its expression correlates with APOBEC3-linked mutational signature burdens in certain cancer types, leading to its initial designation as the predominant APOBEC3 mutator^5,8^. However, multiple recent independent studies instead point to APOBEC3A as the primary driver of APOBEC3 mutagenesis. We demonstrated that APOBEC3A deletion in human cancer cell lines markedly reduces the acquisition of the majority of SBS2/13 mutational burdens, whereas APOBEC3B deletion has a modest effect^4^. Consistent with this, SBS2/13 in tumor genomes are more frequently enriched in the APOBEC3A-preferred YTCA sequence context than in the APOBEC3B-preferred RTCA context (Y/R=pyrimidine/purine bases)^9^. These collective findings, together with the greater intrinsic deaminase potency of APOBEC3A^10^ and additional evidence emerging from a growing number of studies^4,9,11–16^, indicate that APOBEC3A is the most mutagenic APOBEC3 paralog across human cancers. The extent of APOBEC3B contribution may vary across contexts and remains an area of active investigation, challenged by the poor correlation between APOBEC3B expression and SBS2/13 signature burdens^4–7,17,18^.

Importantly, the activity and relative contributions of APOBEC3A and APOBEC3B remain uncertain in cancer types in which SBS2/13 mutations exhibit tumor-to-tumor heterogeneity in RTCA versus YTCA context enrichment, rather than the YTCA dominance observed in most tumor types, and for which genetic delineation has not been performed. This ambiguity is especially pronounced in LUAD suggesting a potentially distinct or comparatively more significant role for APOBEC3B in this malignancy^9,12^. Although APOBEC3-mediated therapy resistance in LUAD has been linked to APOBEC3A and APOBEC3B induction by certain inhibitors of oncogenic signaling pathways, this induction is heterogeneous and incompletely defined across tumors^6,19^. A key unresolved question is whether otherwise prevalent, endogenous misregulation of APOBEC3A and APOBEC3B^4,20^ contributes to mutagenesis in this disease, and with what relative roles—knowledge that is essential for rational therapeutic targeting and that may guide interpretation of therapy-associated effects.

To address this gap, here we resolve the contributions of endogenously misregulated APOBEC3A and APOBEC3B to mutagenesis in human LUAD cells by combining CRISPR–Cas9–mediated deletion of these enzymes with lineage-resolved whole-genome sequencing (WGS). By tracking mutation accumulation across LUAD models that recapitulate the APOBEC3A- and APOBEC3B-associated sequence-context biases observed in tumors, we definitively assign APOBEC3-linked mutational signatures to their enzymatic sources for the first time in this cancer type. Using these genetically defined systems, we further assess the accuracy of commonly used surrogate assays for APOBEC3A and APOBEC3B activities and causally evaluate the proposed link between endogenous APOBEC3 mutagenesis and the indel signature ‘InD9a’^13,21^. Our findings reveal highly heterogeneous contributions of endogenously misregulated APOBEC3A and APOBEC3B to both SBS2/13 and InD9a signature in LUAD and indicate that widely used assays fail to reliably capture their activities in individual tumors, thus highlighting the need for more precise biomarkers to inform therapeutic targeting with APOBEC3 inhibitors currently in development^22–24^.

## Results

### LUAD cell lines recapitulate heterogeneous APOBEC3 mutational sequence contexts in patient tumors

To investigate the primary source of APOBEC3-associated mutational signatures in LUAD, we examined their extended sequence context patterns across 1,686 pan-cancer whole-genomes analyzed by Jalili et al^12^. Consistent with a previous report^9^, LUAD patient tumors displayed substantial heterogeneity in YTCA/RTCA enrichment ratios, with markedly fewer tumors exhibiting strong YTCA bias (**Fig. 1a**). This heterogeneity distinguishes LUAD from cancers such as head and neck and bladder, where YTCA enrichment predominates more uniformly. To determine whether this diversity is recapitulated and could thus be pursued *in vitro*, we analyzed whole-exome sequencing data from 64 LUAD cell lines, confirming that they mirror the YTCA/ RTCA enrichment patterns observed in patient tumors (**Fig. 1a**) and thus provide suitable models to genetically interrogate the contributions of APOBEC3A and APOBEC3B paralogs to mutagenesis.

**Figure 1.**
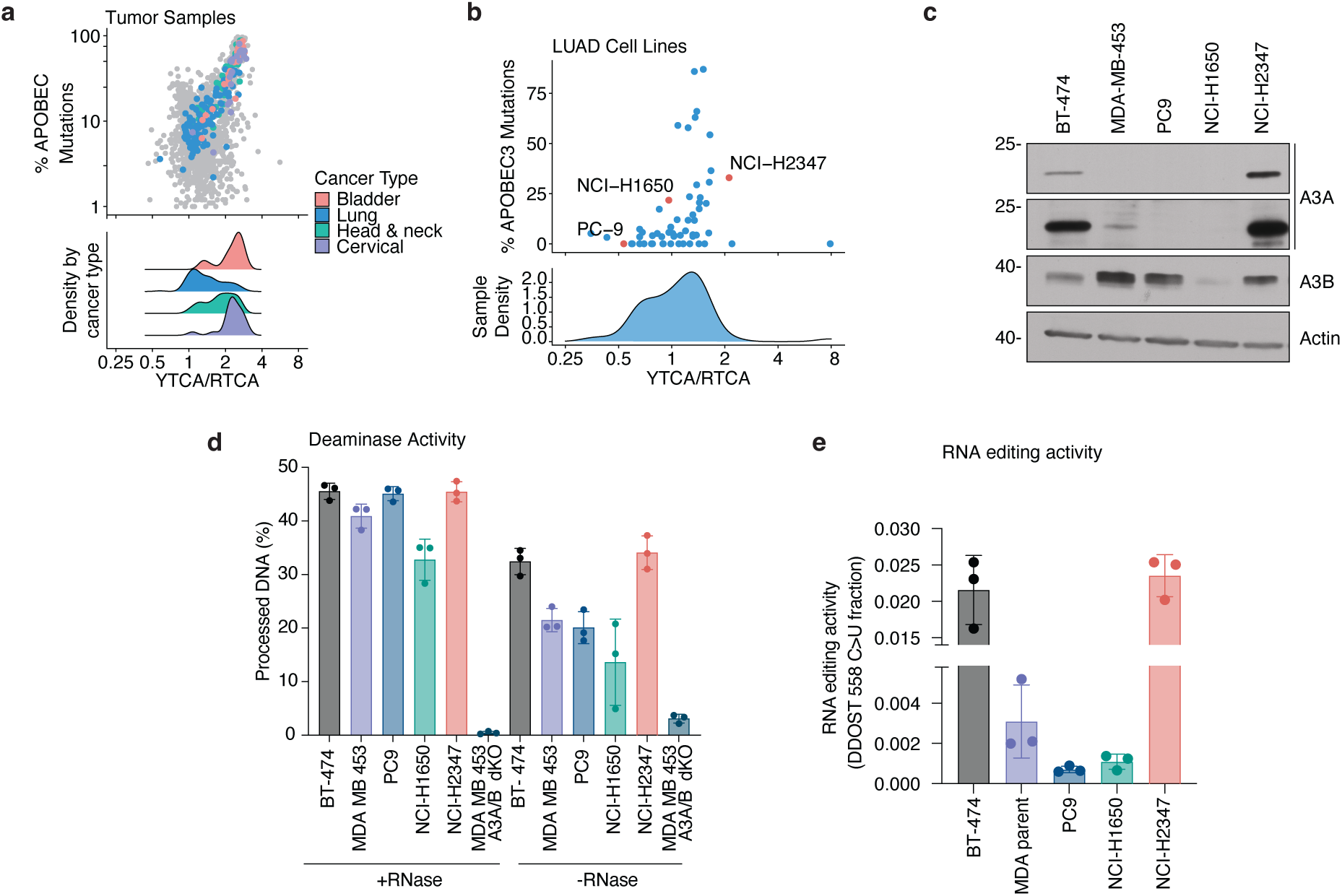
APOBEC3 activity and mutational landscapes in LUAD tumors and cell lines. (a) Enrichment of APOBEC3-associated mutations at YTCA (APOBEC3A-preferred) vs. RTCA (APOBEC3B-preferred) sequence contexts (Y=pyrimidine base; R= purine base; underlined base = APOBEC3-targeted cytosine in TC context). Top: Colored dots indicate individual tumors from color-coded cancer types otherwise exhibiting high SBS2/13 burdens, while grey indicates other cancer types. Y-axis denotes the proportion of mutations attributed to SBS2/13, and X-axis indicates the ratio of mutations occurring in YTCA vs RTCA sequence context. Bottom: Density distributions of sequence context enrichment for indicated cancer types. (b) Top: Enrichment of APOBEC3-associated mutations at YTCA (APOBEC3A-preferred) vs. RTCA (APOBEC3B-preferred) sequence contexts across 64 whole-exome-sequenced LUAD cell lines. The axes are plotted as in (a), highlighting similar YTCA/RTCA enrichment heterogeneity between patient tumors and cell lines. Cell lines selected for further analysis are highlighted in red. Bottom: Density distribution of sequence context enrichment for the 64 LUAD cell lines plotted above. (c) Immunoblot analysis of APOBEC3A (A3A) and APOBEC3B (A3B) protein levels in the indicated breast (BT-474, MDA-MB-453) and LUAD (PC9, NCI-H1650, NCI-H2347) cell lines. Actin is shown as a loading control. (d) In vitro DNA deaminase activity of whole-cell lysates assayed using a linear oligonucleotide substrate in the presence or absence of RNase A. Lysate from an MDA-MB-453 APOBEC3A/APOBEC3B dKO (A3A/A3B dKO) cell line serves as a negative control. Data are mean ± s.d., and points represent three independent biological replicates. (e) Quantification of endogenous APOBEC3A-specific RNA editing activity at the DDOST C558 hotspot. Data are presented as mean ± s.d., points represent independent biological replicates.

From this panel, we selected LUAD cell lines representative of the observed diversity for in-depth analysis, including models widely used to study APOBEC3 mutagenesis. These included NCI-H2347, which exhibits SBS2/13 with cytosine mutations enriched in the APOBEC3A-preferred YTCA contexts; PC9, commonly used in studying APOBEC3 mutagenesis^6,12,19,25,26^, with cytosine mutations skewed towards APOBEC3B-preferred RTCA contexts but lacking detectable SBS2/13, suggesting that these mutations might not in fact reflect APOBEC3 activity; and NCI-H1650, which shows no clear enrichment for SBS2/13 cytosine mutations in either YTCA or RTCA sequence contexts (**Fig. 1b**).

We first characterized these lines using common surrogate assays for APOBEC3 mutagenesis, including measurements of protein, mRNA, and enzymatic activities^22^. Immunoblotting revealed high APOBEC3A protein levels in NCI-H2347, consistent with its YTCA-dominant signature, but no detectable APOBEC3A in PC9 and NCI-H1650 (**Fig. 1c**). APOBEC3B protein was detected at variable levels across all three lines (**Fig. 1c**).

Given the distinct APOBEC3 protein expression patterns across the cell lines, we next measured the APOBEC3 enzymatic activities using an *in vitro* DNA cytosine deaminase assay on whole-cell lysates. Because cellular RNA potently inhibits APOBEC3B but not APOBEC3A^11,12^, assays were performed in the absence of RNase to capture RNA-insensitive activity and in the presence of RNase to unmask the RNA-inhibited activity of RNA-sensitive APOBEC3 deaminases, including APOBEC3B. Under RNase-free conditions, NCI-H2347, characterized by a comparatively higher SBS2/13 mutational burden and detectable expression of both APOBEC3A and APOBEC3B, exhibited the highest deaminase activity on APOBEC3-preferred substrates. In contrast, PC9 and NCI-H1650, which lack detectable *APOBEC3A* expression and exhibit variable APOBEC3B levels, yet show respectively absent or lower SBS2/13 mutational burdens (**Fig. 1b**), displayed lower but detectable deaminase activity (**Fig. 1d**). Upon treatment with RNase, deaminase activity markedly increased across all cell lines, likely reflecting RNA-inhibited APOBEC3B activity (**Fig. 1d**). These results show that deaminase activity broadly tracks with historical SBS2/13 burdens and protein expression in NCI-H2347, but the relationship is less clear in PC9 and NCI-H1650, where detectable enzymatic activity persists despite minimal or absent SBS2/13 signatures.

To specifically probe APOBEC3A’s contribution using an orthogonal approach, we used a highly sensitive RNA editing assay targeting a known hotspot in the *DDOST* transcript, at which APOBEC3A exhibits robust RNA editing whereas APOBEC3B has negligible effects, rendering this assay a specific readout for APOBEC3A activity^12^. NCI-H2347 cells displayed high levels of RNA editing activity (**Fig. 1e**), consistent with their high APOBEC3A protein expression. In contrast, PC9 and NCI-H1650 cells had ∼10-fold lower levels of editing activity (**Fig. 1e**). Thus, unlike the DNA deaminase assay, RNA editing activity correlates well with APOBEC3A protein levels across all three cell lines.

Together, these findings reveal inconsistencies between APOBEC3 mutational signatures, deaminase activity measurements, and protein expression levels. While NCI-H2347 shows concordant results across assays, including high levels of APOBEC3A protein and APOBEC3A-specific RNA editing, accompanied by the SBS2/13 enrichment in APOBEC3A-preferred sequence contexts, PC9 and NCI-H1650 display conflicting genomic and biochemical profiles. Specifically, PC9 exhibits enrichment of cytosine mutations in APOBEC3B-preferred RTCA contexts yet possesses low numbers of SBS2/13 mutations in stock cultures, despite measurable deaminase activity in lysates and maintained APOBEC3B protein expression. This indicates that RTCA context enrichment, commonly interpreted as evidence of APOBEC3B-dominant mutagenesis in tumors, might not accurately reflect ongoing APOBEC3B mutagenic activity. NCI-H1650 further highlights assay discrepancies: it exhibits SBS2/13 without clear enrichment in APOBEC3A- or APOBEC3B-preferred sequence context and displays substantial deaminase activity, both with and without RNase treatment, suggesting both APOBEC3A and APOBEC3B enzymatic activities, despite undetectable APOBEC3A protein and relatively low APOBEC3B protein levels. Therefore, commonly used surrogate measures, including protein levels, mutational sequence context enrichment, and deaminase activity assays, may not reliably infer APOBEC3A and APOBEC3B mutagenic activities. The examined cell line panel represents a fully characterized system to genetically dissect the genuine mutagenic contributions of APOBEC3 paralogs, and the extent to which common surrogate assays correctly infer them.

### Single-cell derived LUAD lineages uncover subclonal APOBEC3 expression that obscures surrogate activity assays

To investigate whether the inconsistent mutational and biochemical profiles in bulk cell lines (**Fig. 1**) reflect subclonal heterogeneity in *APOBEC3A and APOBEC3B* expression, we generated and analyzed a panel of 18 single-cell-derived wild-type (WT) lineages from NCI-H2347, NCI-H1650, and PC9. Immunoblotting revealed that APOBEC3A protein was consistently undetectable in NCI-H1650 lineages and uniformly high in NCI-H2347 lineages, whereas PC9 lineages exhibited striking subclonal heterogeneity. Although APOBEC3A protein was undetectable in the PC9 parental population (**Fig. 1c**), several of its lineages expressed readily detectable APOBEC3A protein (**Fig. 2a**), demonstrating that APOBEC3A-positive subpopulations are masked in bulk analysis. In contrast, APOBEC3B protein levels were relatively uniform across NCI-H1650 and NCI-H2347 clones (**Fig. 2b,c**), while showing a modest increase in variation across PC9 lineages (**Fig. 2a**). To determine whether observed heterogeneity reflected differences in transcription, we quantified *APOBEC3A* and *APOBEC3B* mRNA in examined lineages (**Fig. 2d-f**). qPCR revealed variable mRNA levels among clones that correlated poorly with protein levels, indicating that mRNA abundance does not reliably predict APOBEC3 protein expression^27^. Together, these data demonstrate substantial variability in the expression of APOBEC3 paralogs within highly related, clonally-derived cell line populations and indicate that post-transcriptional regulation, rather than transcription alone, may present a major determinant of APOBEC3 protein abundance.

**Figure 2.**
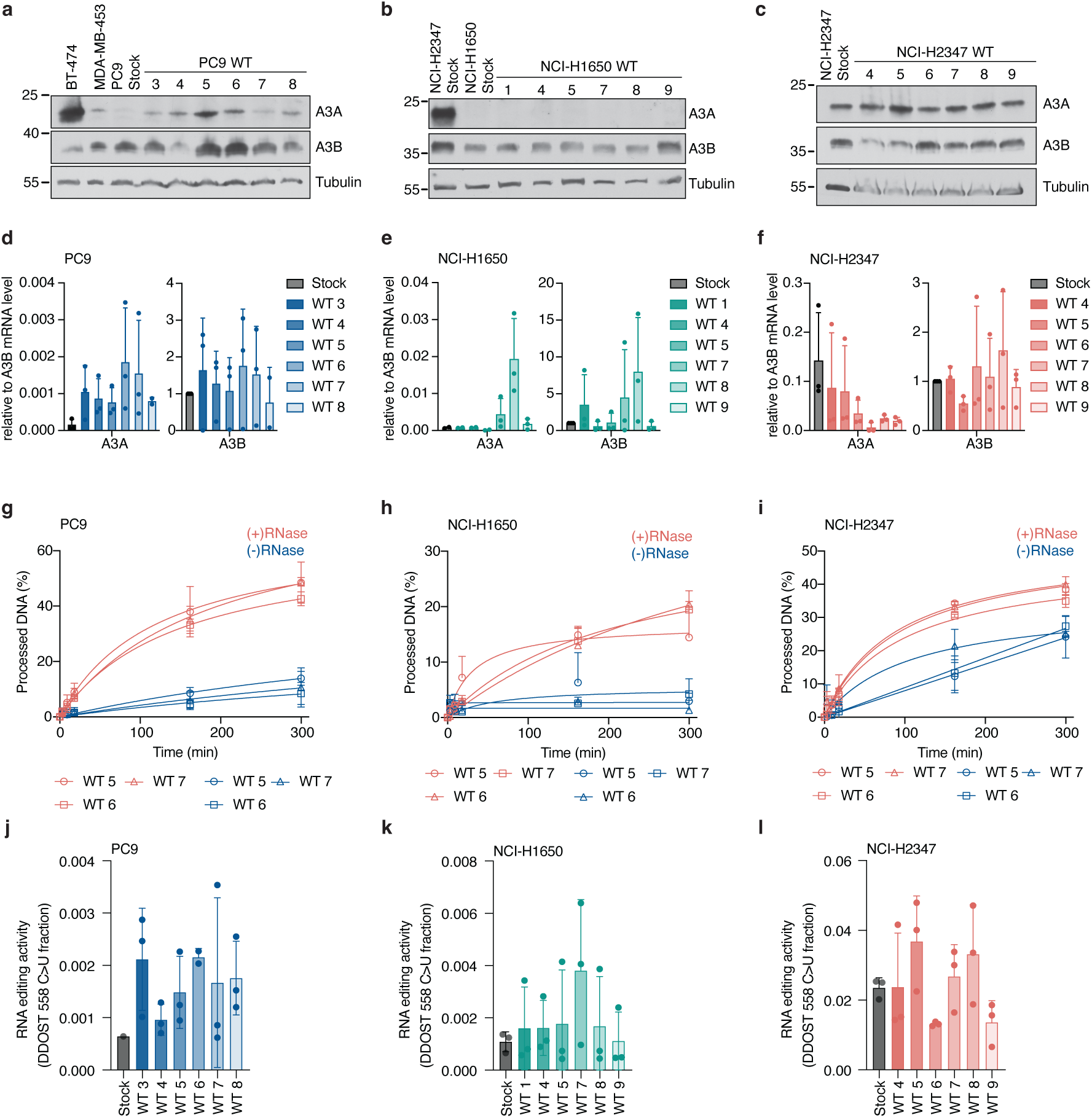
Subclonal heterogeneity of APOBEC3 expression and activity in LUAD cell lines. (a-c) Immunoblot analysis of APOBEC3A (A3A), APOBEC3B (A3B), and loading controls (Actin/Tubulin) in a panel of single-cell-derived wild-type (WT) subclones from (a) PC9, (b) NCI-H1650, and (c) NCI-H2347. Lysates from the corresponding stock cell line population are included for comparison. Lysates from BT-474 and NCI-H2347 samples are included as positive controls for APOBEC3A protein levels. (d-f) qPCR analysis of APOBEC3A (A3A) and APOBEC3B (A3B) mRNA abundance in WT subclones from (d) PC9, (e) NCI-H1650, and (f) NCI-H2347. (g-i) In vitro DNA deaminase activity of whole-cell lysates from WT subclones of (g) PC9, (h) NCI-H1650, and (i) NCI-H2347, measured using a linear oligonucleotide substrate. Samples were assayed in the presence (red traces) or absence (blue traces) of RNase A to distinguish RNase-sensitive and RNase-insensitive deaminase activities. (j-l) Quantification of endogenous APOBEC3A-specific RNA editing activity at the DDOST C558 hotspot in WT subclones from (j) PC9, (k) NCI-H1650, and (l) NCI-H2347. All bar graphs (d-l) show mean +/- s.d. for three independent biological replicates. Statistical analyses in (j-l) were performed using an ordinary one-way ANOVA with Tukey’s multiple-comparisons test.

We next tested whether the expression heterogeneity translated to functional differences in enzymatic activity. Deaminase assays across the characterized lineages revealed the expected distinction between RNase-sensitive and RNase-insensitive activities^11^, reflecting APOBEC3B and APOBEC3A, respectively (**Fig. 2g-i**). However, within each condition, the ranges in protein and mRNA levels did not translate to corresponding differences in deaminase activity. To specifically assess APOBEC3A activity, we quantified RNA editing of the *DDOST* transcript across the lineage panel. Editing levels varied approximately two-fold among lineages within each cell line (**Fig. 2j-l**), but did not track with APOBEC3A protein abundance, possibly reflecting the limited signal-to-noise ratio and dynamic range of this assay for detecting modest differences in activity. Across PC9 and NCI-H1650 lineages, RNA editing was similarly low (**Fig. 2j, k**), whereas those from NCI-H2347 exhibited substantially higher activity (**Fig. 2l)**, consistent with the higher APOBEC3A protein levels (**Fig. 2c**) in this cell line.

Together, these results demonstrate that APOBEC3A expression can be highly subclonal and heterogeneous across lineages, such that APOBEC3A activity may appear absent at the population level despite being active in discrete cancer cell subsets. APOBEC3B expression likewise varies across lineages, further compounding inference from bulk measurements. These findings expose a fundamental limitation of bulk and surrogate assays for inferring APOBEC3 mutagenic activity in cancer: subclonal heterogeneity can mask both the presence and relative contributions of individual APOBEC3 enzymes within tumor populations. Moreover, expression-based readouts and enzymatic assays are constrained by limited dynamic range, obscuring modest but biologically meaningful differences in activity. Together, these limitations necessitate genetically defined knockout models to unambiguously resolve the sources of APOBEC3-driven mutagenesis.

### Genetically dissecting contributions of endogenous APOBEC3A and APOBEC3B to genome-wide mutagenesis in LUAD

To overcome the limitations of surrogate assays and definitively determine the contributions of endogenous APOBEC3 paralogs to mutagenesis in characterized LUAD cell lines, we used CRISPR-Cas9 to delete APOBEC3A and APOBEC3B individually or in combination. We then applied our established workflow for tracking *de novo* mutations across WT and knockout (KO) single-cell-derived lineages (**Fig. 3a**)^4,20^.

**Figure 3.**
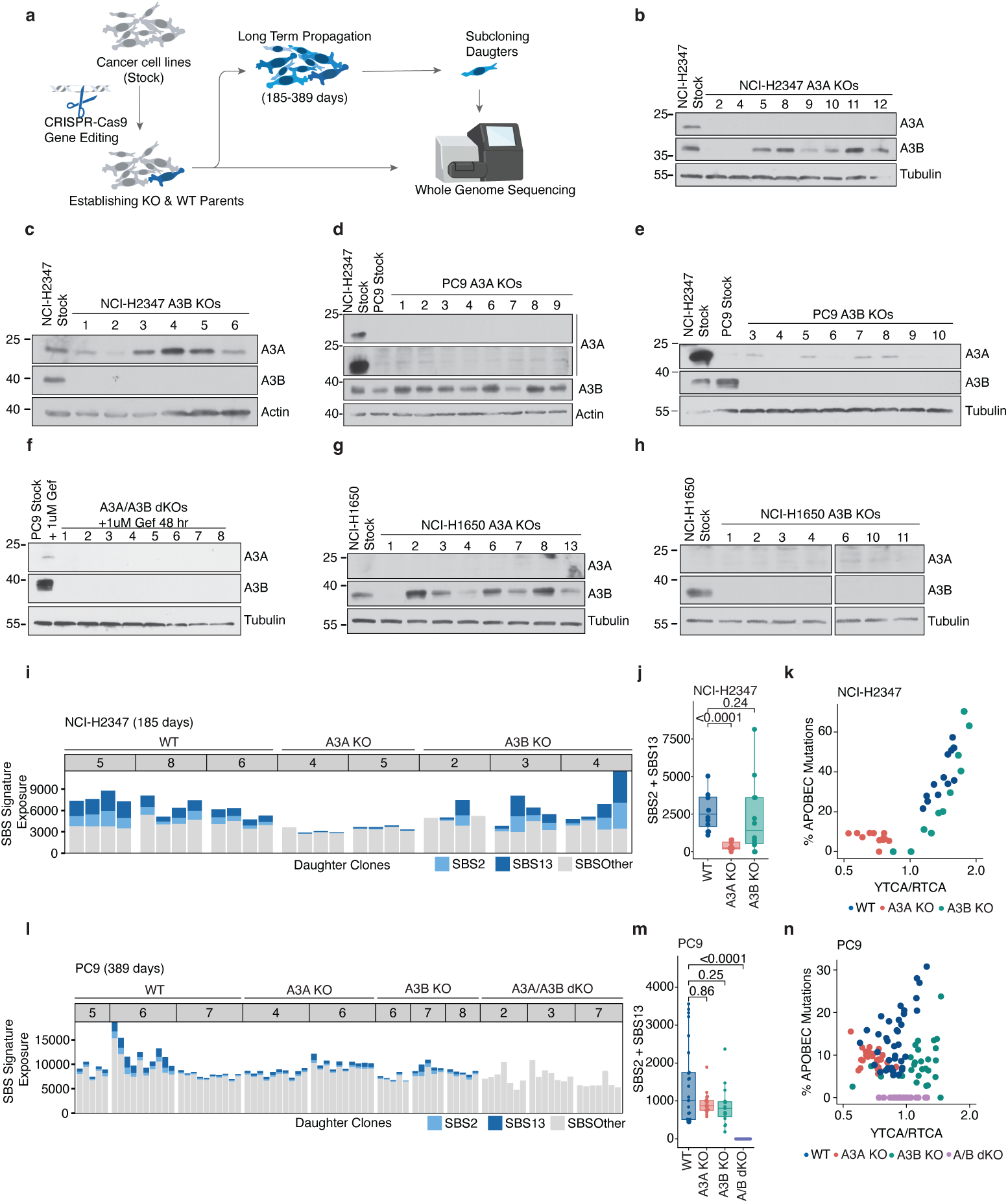
APOBEC3 mutagenic activities in LUAD are highly heterogeneous and context-dependent. (a) Schematic of the longitudinal mutational analysis workflow used to quantify *de novo* mutation accumulation across genetically defined lineages. (b-h) Immunoblot validation of APOBEC3 KO clones. (b–h) Immunoblot validation of APOBEC3 knockout (KO) clones showing APOBEC3A (A3A) and APOBEC3B (A3B) protein levels in NCI-H2347, PC9, and NCI-H1650 APOBEC3A KO, APOBEC3B KO, and APOBEC3A/APOBEC3B double-KO (dKO) clones. Tubulin and actin are shown as loading controls. Lysates from NCI-H2347 stock cells (untreated) or PC9 stock cells treated with gefitinib^19^ are included as reference samples with detectable APOBEC3A protein expression. (i) *De novo* SBS mutations accumulated genome-wide in individual NCI-H2347 daughter clones (WT, APOBEC3A KO/A3A KO, APOBEC3B KO/. A3B KO) following >150 days of propagation. Colors indicate inferred mutational etiologies assigned by mutational signature analysis. (j) Grouped analysis of SBS2/13 mutation accumulation in NCI-H2347 clones. Box plots show the median (center line), 25th and 75th percentiles (box limits), and whiskers extending to 1.5 x interquartile range. Each dot represents the combined SBS2/13 mutation burdens in each individual daughter clone. P-values shown were calculated using a two-tailed Mann-Whitney U test comparing SBS2+SBS13 burdens in each knockout genotype to the wild-type (WT) of the same cell line and time point. (k) Enrichment of of *de novo* APOBEC3-associated mutations from NCI-H2347 daughter clones in APOBEC3B-preferred RTCA vs APOBEC3A-preferred YTCA sequence contexts. Each point represents a single clone, colored by genotype. (l) *De novo* SBS mutations accumulated in individual PC9 daughter clones (WT, A3A KO, A3B KO, A3A/A3B dKO) propagated for 389 days. (m) Grouped analysis of SBS2/13mutation accumulation in PC9 clones. Data presentation and statistical testing as in (j). (n) Enrichment of *de novo* APOBEC3-associated mutations from PC9 daughter clones in APOBEC3B-preferred RTCA vs APOBEC3A-preferred YTCA sequence contexts. Each point represents one clone, colored by genotype.

Knockout efficiency was validated across derived lineages at genetic, protein, and functional levels. Loss of APOBEC3A and APOBEC3B expression was confirmed by comprehensive immunoblotting, RT-qPCR, and targeted Sanger sequencing (**Fig. 3b-h; Supplementary Fig. 2, Supplementary Table 1**). Importantly, qPCR analysis demonstrated that expression of other *APOBEC3* family members was not systematically altered by the genetic perturbations (**Supplementary Fig. 2**). Functional validation corroborated complete enzymatic inactivation: DNA deaminase assays showed that *APOBEC3B* deletion eliminated RNase-sensitive deaminase activity, whereas *APOBEC3A* deletion specifically reduced RNase-insensitive activity, particularly in NCI-H2347 (**Supplementary Fig. 3a-j**), consistent with the known biochemical properties of APOBEC3B and APOBEC3A. Finally, *DDOST* RNA editing assays demonstrated a nearly complete loss of APOBEC3A-specific activity in the *APOBEC3A* KO lineages, confirming functional inactivation at the cellular level (**Supplementary Fig. 3k-m**).

Obtained single-cell–derived lineages of WT, *APOBEC3A* KO, *APOBEC3B* KO, and *APOBEC3A*/*APOBEC3B* double KO (dKO) cells were independently subjected to long-term cultivation to enable mutation acquisition. Lineages from NCI-H2347 and NCI-H1650 were propagated for >150 days, whereas PC9 lineages were propagated for an extended duration with samples collected at two time points (178 and 389 days). Given the low endogenous mutation load in PC9, this extended propagation was necessary to accumulate sufficient mutations for analysis. All selected lineages exhibited proliferation rates similar to the parental stock, ensuring that differences in mutation accumulation are not attributable to growth defects (**Supplementary Fig. 1**).

After these propagation periods, each parental lineage was subcloned into multiple single-cell-derived daughter clones for genomic analysis. We performed 30× WGS on DNA from these daughter clones as well as on DNA from the corresponding parental lineage obtained shortly after isolation of the founding cell. Because 30× WGS of clonal populations primarily detects clonal mutations, sequencing of the parental DNA largely captures pre-existing mutations present in each examined lineage at the start of the experiment. In contrast, sequencing of daughter clones captures these mutations together with those acquired during propagation up to the point at which individual daughter cells were isolated at the end of the propagation window. WGS data obtained from the parental lineages was leveraged to additionally validate successful knockouts of the targeted proteins. Mutations acquired *de novo* during the propagation windows were defined as variants present in individual daughter clones but absent from their respective parental lineages, and were then used to extract and assign mutational signatures, identifying APOBEC3-associated SBS2/13 (Methods, **Supplementary Fig. 4-8**). By comparing *de novo* SBS2/13 accumulation across genotypes, we could causally attribute genome-wide mutagenesis to specific APOBEC3 enzymes.

Analyses of mutations acquired across lineages from NCI-H2347, which exhibits strong APOBEC3A-preferred mutational patterns, revealed that WT daughter lineages accumulate a high burden of *de novo* SBS2/13 mutations (**Fig. 3i-j, Supplementary Fig. 4,11**). SBS2/13 burdens were severely reduced, though not completely eliminated, in the *APOBEC3A* KO lineages (p < 0.0001) (**Fig. 3i,j, Supplementary Fig. 4,11**). In contrast, knockout of *APOBEC3B* had no significant effect on the rate of SBS2/13 accumulation compared to WT lineages (**Fig. 3i-j, Supplementary Fig. 4,11**). Analysis of the mutational extended sequence contexts confirmed that *de novo* mutations in WT lineages were strongly enriched toward the APOBEC3A-preferred YTCA motifs. The *APOBEC3B* KO lineages retained this YTCA bias, while the mutations acquired in *APOBEC3A* KO lineages showed a dramatic reduction in YTCA-context enrichment with minimal burdens of residual SBS2/13 mutations (**Fig. 3k**). These results establish APOBEC3A as the primary driver of mutagenesis in NCI-H2347 cells, as predicted by the surrogate assays, with APOBEC3B contributing negligibly. This APOBEC3A dominance in models with SBS2/13 enriched in YTCA sequence contexts is consistent with our previous findings in breast cancer and lymphoma cell lines^4^.

In PC9 cells, which exhibit low SBS2/13 burdens enriched in RTCA contexts and subclonal APOBEC3A expression, deletion of either *APOBEC3A* or *APOBEC3B* reduced the variance in SBS2/13 mutations but did not result in a statistically significant reduction in these mutations relative to WT lineages (APOBEC3A-KO *p = 0.86,* APOBEC3B-KO *p = 0.25*) (**Fig. 3l-m, Supplementary Fig. 5,11**). Acquisition of SBS2/13 mutations was completely abolished in the *APOBEC3A/APOBEC3B* dKO lineages (*p < 0.0001*) (**Fig. 3l-m, Supplementary Fig. 5, 11**). Extended sequence-context analysis revealed that d*e novo* mutations in *APOBEC3A*-knockout lineages were strongly RTCA-enriched, consistent with residual APOBEC3B activity, whereas mutations in *APOBEC3B*-knockout lineages were YTCA-enriched, reflecting remaining APOBEC3A activity. dKO lineages lacked the characteristic YTCA or RTCA context enrichment observed in WT and single-knockout lineages (**Fig. 3n**). Together, these data demonstrate that APOBEC3A and APOBEC3B are comparably active mutators in PC9 cells. Notably, although this dual contribution is evident from subclonal APOBEC3A and APOBEC3B expression across the examined WT single-cell-derived lineages, it would not have been inferred from surrogate assays performed on the cell line stock population, which exhibited undetectable bulk APOBEC3A protein, low *DDOST* RNA editing activity, and APOBEC3B-preferred, RTCA-biased mutational patterns. These findings underscore the value of causal genetic approaches for definitive understanding of APOBEC3 mutagenic activities.

The complete loss of SBS2/13 acquisition in PC9 *APOBEC3A*/*APOBEC3B* dKO lineages was notable given the persistence of residual deaminase activity *in vitro* (**Supplementary Fig. 3j**). This activity was specific to TCA-containing substrates and not observed on a disfavored ACA motif (**Supplementary Fig. 9a**), indicating involvement of other APOBEC3 paralogs. Consistent with this, PC9 dKO cells expressed *APOBEC3C*, *APOBEC3F*, and *APOBEC3H* transcripts (**Supplementary Fig. 2c**). Targeted knockdown of these paralogs revealed that only APOBEC3F contributed to the residual TCA deaminase activity (**Supplementary Fig. 9b**). However, APOBEC3F-GFP localized exclusively to the cytoplasm and was excluded from chromatin at all cell cycle stages (**Supplementary Fig. 9c**), consistent with prior reports^28^. Thus, despite measurable *in vitro* activity, APOBEC3F is highly unlikely to contribute to genomic mutagenesis in PC9 cells. These results underscore the ambiguities inherent in bulk surrogate assays, as the APOBEC3F-driven *in vitro* activity detected in lysates is highly unlikely to translate to genuine genomic mutagenesis.

In contrast to NCI-H2347 and PC9, most NCI-H1650 WT lineages did not acquire SBS2/13 mutations during propagation (**Supplementary Fig. 6, 10a,b, 11b**), despite this cell line exhibiting an intermediate historical SBS2/13 mutational burden with mixed YTCA/RTCA enrichment (**Fig. 1b; Supplementary Fig. 6**). This contrasts with PC9, which has a lower historic SBS2/13 yet has accumulated substantially more de novo mutations during propagation (**Fig. 1b; Supplementary Fig. 5**). Notably, SBS2/13 mutations were detected in a single NCI-H1650 WT lineage, where cytosine mutations were enriched in APOBEC3A-preferred YTCA contexts (**Supplementary Fig. 10a,c**). APOBEC3A and APOBEC3B protein levels were not elevated in this lineage relative to others (**Supplementary Fig. 10d**), suggesting that these mutations arose from a rare, episodic burst of APOBEC3A activity^4,20^. Together, these findings indicate that APOBEC3A mutagenic bursts may occur infrequently in NCI-H1650 cells and reinforce that historical SBS2/13 burdens do not necessarily reflect ongoing APOBEC3 mutagenic activity^20^.

Analyses of other signatures confirmed that, as expected, all lineages steadily accumulated background mutations from signatures attributed to aging and cell culture (**Supplementary Fig. 11a-c**). Interestingly, we found a consistent contribution from SBS3, a mutational signature associated with homologous recombination (HR) deficiency, in PC9 lineages, corroborated by the presence of the HR deficiency-associated indel signature InD6 (**Supplementary Fig. 11d**)^29^. To our knowledge, this represents the first indication of a HR deficiency in this commonly used cell line, a finding that may warrant consideration in future studies using this model.

### APOBEC3A and APOBEC3B contributions to clustered mutations in LUAD

In addition to dispersed genome-wide SBS2/13 mutations, APOBEC3 activity also generates characteristic localized hypermutation clusters—*omikli* (medium-density clusters) and *kataegis* (high-density foci)^30–33^. To determine whether the APOBEC3A-versus-APOBEC3B hierarchy we identified for dispersed mutations also extends to these clustered mutational events, we identified *de novo omikli* and *kataegis* events (Methods) in our cell line cohorts.

In NCI-H2347 lineages, acquisition of both *omikli* and *kataegis* was significantly reduced in *APOBEC3A* KO lineages at 148 days (*omikli: p*=0.019; kataegis: *p*=0.037), with no significant effects observed in *APOBEC3B* KOs (**Fig. 4a,b**). This pattern mirrors that seen for dispersed SBS2/13 mutations, establishing APOBEC3A as the sole driver of both dispersed and clustered mutagenesis in this line. In contrast, PC9 lineages at 389 days showed significant decreases in clustered mutagenesis in APOBEC3B KOs (*omikli:* p < 0.001, *kataegis:* p < 0.05) and dKOs (*omikli:* p < 0.0001, *kataegis:* p < 0.05), and no significant effects in APOBEC3A KOs (**Fig. 4c, d**). NCI-H1650 lineages showed few clustered events and no detectable effects of APOBEC3 knockouts, consistent with minimal APOBEC3 activity (**Fig. 4e,f**). Collectively, these analyses demonstrate that the context-dependent hierarchy of APOBEC3A and APOBEC3B governs both dispersed and clustered hypermutational processes.

**Figure 4.**
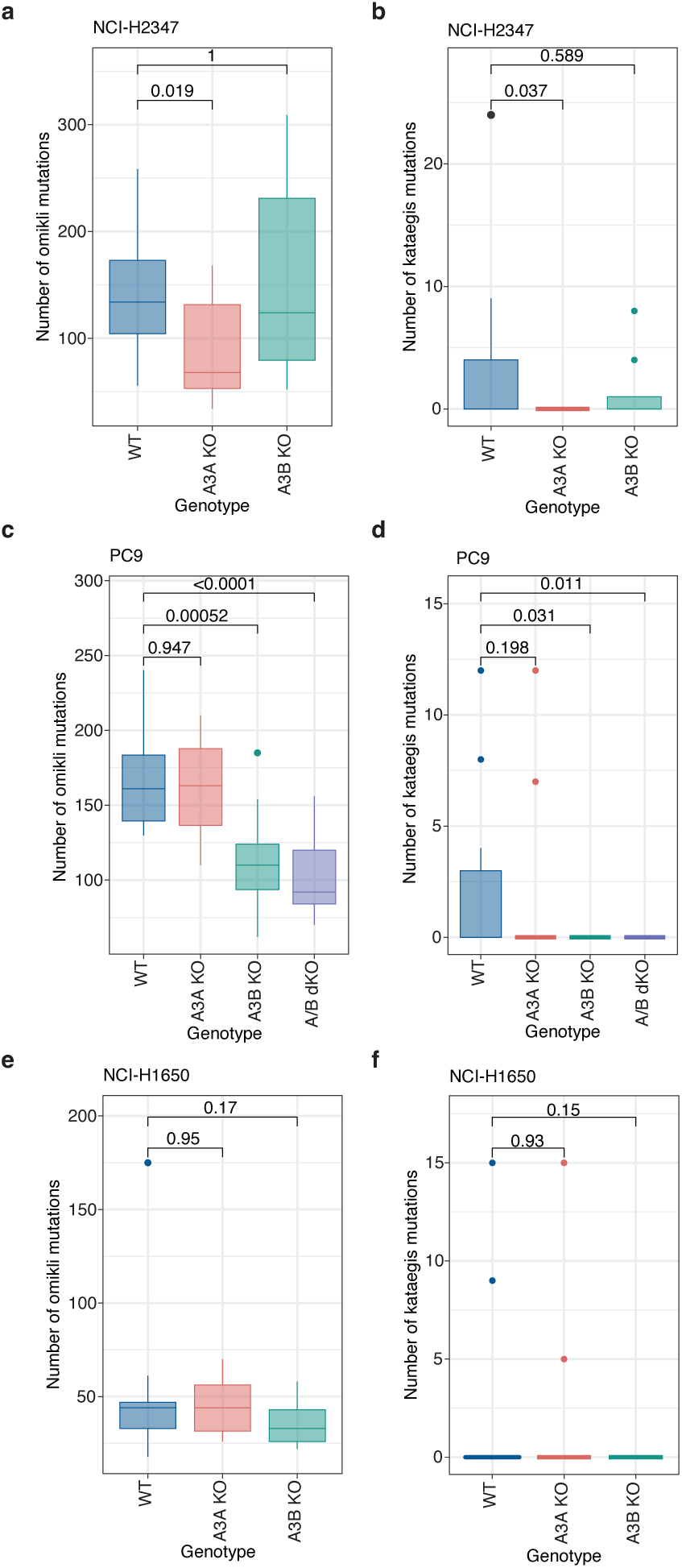
The APOBEC3 mutagenic hierarchy extends to clustered mutational events. (a, c, e) Quantification of *de novo omikli* mutation clusters in daughter clones of the indicated genotypes (a) NCI-H2347 (185 days), (c) PC9 (389 days), and (e) NCI-H1650 (160 days). (b,d,f) Quantification of *de novo kataegis* mutation foci accumulated in daughter clones of the indicated genotypes (b) NCI-H2347 (185 days), (d) PC9 (389 days), and (f) NCI-H1650 (160 days). Panels are ordered as in (a). Box plots show the median (center line), 25th and 75th percentiles (box limits), and whiskers extending to 1.5 x interquartile range. Each dot represents clustered mutational counts in each individual daughter clone. P-values shown were calculated using a two-tailed Mann-Whitney U test comparing clustered mutation counts in each knockout genotype to the wild-type (WT) of the same cell line and time point.

### APOBEC3 deaminases drive the InD9a indel signature

APOBEC3 enzymes are also linked to the generation of indel mutations. Initial evidence came from ectopic expression of APOBEC3A in DT40 chicken cells, which revealed a signature dominated by ’T’ base insertions and 1 bp ’C’ deletions at TCA and TCT motifs^13,21^. This signature correlated with SBS2/13 mutational burdens across examined tumor types and, among census COSMIC indel signatures, was most similar to signature ID9, which is prevalent across cancers. This led to the hypothesis that ID9 represents an APOBEC3-driven mutational signature, although with a discrepancy of ID9 lacking the T insertions observed in the DT40 model^3,13^. Subsequent reclassification of indels incorporating extended sequence contexts resolved this APOBEC3-associated indel signal into the signature ‘InD9a’^21^, refining the broader, context-agnostic ID9 signature into a pattern specifically enriched for 1-bp C deletions at TCA and TCT motifs, characteristic of APOBEC3-associated SBS2/13, frequently occurring within short poly-T tracts. Together, these observations led to a mechanistic model in which APOBEC3-mediated C-to-U deamination at TC motifs within short repetitive T tracts is followed by uracil excision by UNG2, generating an abasic site that promotes template-strand slippage and results in cytosine deletions^13,21^. However, a direct causal link between endogenous human APOBEC3 deaminases and the InD9a indel signature has not been established; the relative contributions of individual APOBEC3 paralogs remain unresolved, and support for this model has relied largely on indirect evidence.

To fully characterize APOBEC3-related indels in our dataset, and to increase power for signature extraction while extending insights across additional cancer types, we extracted indel signatures from a combined dataset comprising *de novo* mutations identified in the cell lines examined here together with previously generated data from WT, *APOBEC3A*-KO, *APOBEC3B*-KO, *APOBEC3A*/*APOBEC3B*-dKO, UNG2-GFP overexpression, and *UNG2*-KO breast (BT-474, MDA-MB-453) and lymphoma (JSC-1, BC-1) cell lines in which we previously characterized SBS2/13 acquisition^4^.

Among others, an indel signature, termed ‘CH89B’, was extracted and resolved into tumor-derived signatures^21^ (Methods), identifying the contribution of ID9a and thereby enabling its quantification across genotypes (**Supplementary Fig. 12a, 12b**). In the NCI-H2347 (**Fig. 5a**) and MDA-MB-453 (**Fig. 5b**) lineages, InD9a acquisition showed a significant dependence on APOBEC3A (both *p* < 0.01), and a trend towards reduced InD9a acquisition in BT-474 *APOBEC3A* KO clones (*p =* 0.072). In PC9 (**Fig. 5a**), InD9a acquisition was significantly reduced in *APOBEC3A*/*APOBEC3B* dKO clones (*p < 0.05* for PC9). The lack of individual knockout effects in the PC9 and BT-474 lines likely reflects lower overall ID9a burdens, limiting power to resolve single-enzyme effects, as well as previously observed heterogeneity of APOBEC3 mutational activity in BT-474 WT clones^4^. In contrast, no significant changes in ID9a acquisition were observed in *APOBEC3B* knockout clones across cell lines. NCI-H1650, BC-1, and JSC-1 lineages showed no appreciable variation in ID9a acquisition, consistent with their overall low InD9a mutational burdens (**Supplementary Fig. 12c**).

**Figure 5.**
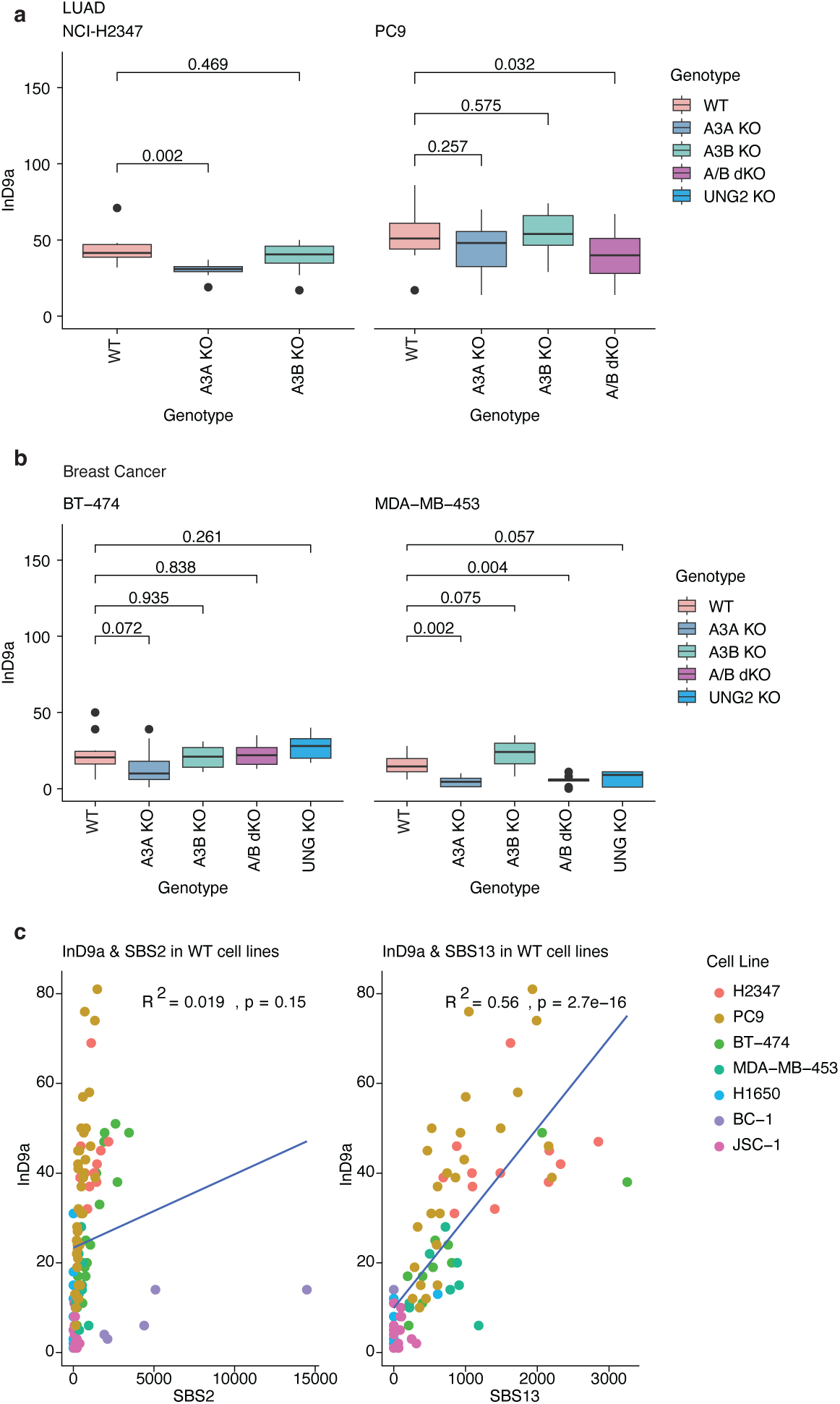
Endogenous APOBEC3 deaminases induce indel signature InD9a found in tumors. (a) Quantification of *de novo* InD9a mutation accumulation in daughter clones from: NCI-H2347 (148 days) and PC9 (389 days) (b) Quantification of *de novo* InD9a mutation accumulation in breast cancer daughter clones from: MDA-MB-453 and BT-474. Box plots show the median (center line), 25th and 75th percentiles (box limits), and whiskers extending to 1.5 times x interquartile range. Dots indicate outlier clones with InD9a mutation burdens beyond the whisker range. P-values shown were calculated using a two-tailed Mann-Whitney U test comparing each genotype to its respective wild-type (WT) control. (c) Correlation of *de novo* acquired InD9a mutational burden with *de novo* acquired SBS2 and SBS13 mutation burdens. Each point represents an individual wild-type (WT) daughter clone across 18 breast, 45 lung, and 21 lymphoma clones from NCI-H1650, NCI-H2347, PC9, BT-474, BC-1, MDA-MB-453, and JSC-1 cell lines. The R2 and P-values from simple linear regressions are shown.

To investigate the proposed model of UNG2-dependent etiology of ID9a, we analyzed ID9a burdens across available clones from cell lines with genetically modified UNG2. Consistent with the model, we found a trend towards reduced ID9a burden in *UNG2*-KO MDA-MB-453 clones (*p* = 0.057) (**Fig. 5b**). In contrast, *UNG2*-KO BT-474 clones and UNG2-GFP knock-in BC-1 clones showed no significant effects on InD9a acquisition (**Supplementary Fig. 12c, Fig. 5b**). The lack of effect in BT-474 likely reflects the substantial inter-clonal heterogeneity in APOBEC3 mutational activity previously characterized in this line^4,20^, which limits statistical power to detect modest reductions. In BC-1, which is otherwise UNG2-deficient^20^, UNG2-GFP overexpression failed to increase InD9a despite partially restoring SBS13^4^, likely reflecting the modest SBS13 levels achieved in this system.

Finally, to further investigate the relationship between InD9a and APOBEC3-driven SBS2/13 signature mutations, we analyzed correlations between *de novo* InD9a and SBS2/13 burdens across the examined WT lineages. While prior studies reported correlations between ID9/InD9a and cumulative SBS2/13 burdens in tumors^13,21^, analysis of SBS2 and SBS13 individually revealed that this relationship is driven almost exclusively by SBS13 (**Fig. 5c**, R^2^ = 0.56, p = 2.7e-16), otherwise dependent on UNG2^4^, with no detectable correlation with SBS2 (**Fig 5c**, *R^2^ = 0.019, p = 0.15*). Together with the APOBEC3 and UNG2 genetic perturbation data, these results support a model in which ID9a arises specifically from APOBEC3-driven, UNG2-dependent processing events underlying SBS13.

Collectively, these data provide the first causal evidence that endogenous APOBEC3A drives InD9a generation in human cancer cells. While a minor contribution from APOBEC3B cannot be excluded, any such effects were below the limit of detection, likely due to substantially lower overall burdens of ID9a compared to SBS2/13. The strong correlation between InD9a and SBS13, but not SBS2, further supports the model that both mutation types arise through the same abasic site-dependent pathway following UNG2-mediated excision, rather than through direct replication across uracil lesions.

## Discussion

Despite the clinical importance of understanding APOBEC3-driven mutagenesis in LUAD, particularly given its role in tumor evolution and therapy resistance^4–7^, the relative contributions of endogenous APOBEC3A and APOBEC3B to mutagenesis have remained uncertain. To resolve this question, we applied a rigorous genetic and longitudinal WGS approach to LUAD cell lines representing the range of YTCA/RTCA enrichment patterns observed in patient tumors.

Our findings provide a framework for interpreting APOBEC3 paralog contributions to mutagenesis in cancer types with tumors with highly heterogeneous YTCA/RTCA enrichment patterns. In YTCA-enriched tumors, as modeled by NCI-H2347, APOBEC3A is likely the sole driver of mutagenesis, with APOBEC3B contributing negligibly despite detectable expression.

In contrast, RTCA-enriched tumors may reflect contexts where both APOBEC3A and APOBEC3B can be equally active and contribute to the mutational burden, as observed in PC9, consistent with recent reports suggesting combinatorial contributions of these enzymes in other cancer types^34^. Importantly, even tumors with minimal APOBEC3 mutational signatures may harbor ongoing, likely episodic^20^, APOBEC3A activity affecting rare clonal fractions, as exemplified by NCI-H1650. While the minimal APOBEC3B mutagenesis in NCI-H1650 likely reflects lower protein abundance, the robust combined APOBEC3A and APOBEC3B activity in PC9 coincides with a distinct homologous recombination-deficient background (SBS3, ID6). This genomic instability may further potentiate APOBEC3B mutagenesis in PC9 by generating the single-stranded DNA substrates required for enzymatic activity^35^. This context-dependent variation underscores that accurate prediction of therapeutic vulnerabilities will require direct assessment of enzyme-specific activities rather than reliance on historical mutational patterns alone.

A key finding of our study is the unreliability of standard surrogate assays for predicting APOBEC3 mutational activity, which helps explain the long-standing ambiguity in the field. We demonstrate that APOBEC3A protein expression, in particular, is highly dynamic and clonally heterogeneous (**Fig. 2**). The highly subclonal nature of APOBEC3A expression confounds efforts to link bulk assays to mutagenesis, rendering population measurements poor predictors. A negative immunoblot result, as we observed in the parental PC9 line, may simply obscure a rare but highly mutagenic subpopulation. While other factors such as immune cell infiltrates with high APOBEC3 expression can also complicate bulk tissue measurements^22,36^, our findings reveal that subclonal heterogeneity within tumor cell populations themselves represents an additional critical limitation. This unreliability extends to functional assays as well. We found that even the highly specific DDOST RNA editing assay^12^ failed to correlate with actual mutagenic output across our lineage panel (**Fig. 2j-l**, **Fig. 3i-n**), despite its sensitivity for detecting APOBEC3A activity. Furthermore, we demonstrate that deaminase activity in cell lysates can be completely decoupled from genomic mutagenesis. We found that residual, TCA-specific deaminase activity in PC9 dKO lysates (**Ext. Data Fig. 9**) was attributable to APOBEC3F. While enzymatically active in lysates, APOBEC3F is strictly cytosolic and thus is highly unlikely to contribute to the nuclear mutational landscape. Together, these data show that surrogate assays, from immunoblots to functional readouts, can be misleading, and that only direct genetic dissection combined with longitudinal sequencing can definitively assign mutational causality.

Beyond the APOBEC3A/APOBEC3B hierarchy, our work also refines the understanding of APOBEC3-driven indel mutagenesis. We genetically confirmed that the InD9a signature, 1bp ’C’ deletions at TCA/TCT motifs^21^, is driven by the same APOBEC3 enzymes responsible for substitutions (**Fig. 5**). Our analysis also revealed a highly specific correlation between InD9a and SBS13 (C>G, C>A), but notably not SBS2 (C>T) (**Fig. 5c**). This selective correlation points to a shared mechanistic origin specifically through the transversion pathway. The prevailing model posits that C>T transitions (SBS2) arise from replication across the initial C>U lesion^4,13,37,38^. In contrast, both C>G transversions (SBS13) and InD9a deletions are thought to arise after the uracil is excised by UNG, creating an abasic site^4,13,37,38^. The correlation between InD9a and SBS13 supports this model, suggesting that InD9a and SBS13 are ’sister’ outcomes of the same abasic-site processing pathway, whereas SBS2 represents a distinct, parallel pathway. Our genetic knockout experiments provide the first direct causal evidence that both APOBEC3A and APOBEC3B generate indel mutations through this mechanism.

Importantly, we confirm that the context-dependent genetic hierarchies we defined here extend beyond dispersed SBS2/13 and InD9a mutations to also include clustered *omikli* and *kataegis* events (**Fig. 4**). Together, our findings provide a foundation for developing enzyme-specific therapeutic strategies and biomarkers. Finally, our findings reinforce the fundamentally episodic nature of APOBEC3 mutagenesis^20^. The NCI-H1650 cell line, which showed minimal activity except for a singular burst of APOBEC3A-like mutations in one clone (**Supplementary Fig. 10**), exemplifies this. This stochastic activity, coupled with the subclonal heterogeneity in APOBEC3A expression we identified (**Fig. 2**), suggests that APOBEC3A is not constitutively expressed. Instead, our data are consistent with a model of stochastic, subclonal induction of APOBEC3A expression, which in turn drives transient bursts of mutagenesis. Understanding the triggers for this potent transcriptional induction, and its full range of cellular consequences, is a critical next step.

## Methods

### Cell Culture

NCI-H1650, NCI-H2347, and PC9 cell lines were acquired from cryopreserved aliquots of cell lines sourced previously from collaborators or public repositories and extensively characterized as part of the Genomics of Drug Sensitivity in Cancer (GDSC) and COSMIC Cell Line projects^20,39,40^. MDA-MB-453 and BT-474 cell lines were sourced as described previously^4^. Individual cell lines and derived clones were genotyped to confirm their identities^4^. All cell lines were mycoplasma negative (LookOut Mycoplasma PCR Detection Kit; Sigma Aldrich MP0035-1KT). MDA-MB-453, NCI-H1650, and NCI-H2347 were grown in RPMI medium supplemented with 10% fetal bovine serum (FBS) and 1% penicillin-streptomycin. BT-474 and PC9 cells were grown in RPMI medium supplemented with 10% FBS, 1% penicillin–streptomycin, 1% sodium pyruvate, and 1% glucose. Unless otherwise noted, all media and supplements were supplied by the MSKCC Media Preparation core facility.

### Generation and validation of knockout and overexpression cell lines

For knockout protocols, cells (2x 10^6^ per 10 cm plate) were transfected using Lipofectamine 3000 Reagent (Invitrogen; L3000015) according to the manufacturer’s protocol. For each transfection, 10 µg of the pU6-sgRNA_CBh-Cas9-T2A-mCherry plasmid, encoding guides targeting either APOBEC3A or APOBEC3B (**Supplementary Table 2**), was used. Double knockout (dKO) clones were established by transfecting validated *APOBEC3B* KO clones with the *APOBEC3A* KO plasmid. Twenty-four hours post-transfection, mCherry-positive cells were single-cell sorted by FACS using the FACSAria system (BD Biosciences) into 96-well plates. APOBEC3F-GFP cells were generated using a piggyBac transposase system. 2.5-3×10^6^ cells were plated in a 10cm plate. 24 hours later, they were transfected using Lipofectamine 3000 Reagent according to the manufacturer’s protocol. The cells received 5 μg of an HP138-APOBEC3F-GFP and 5 μg of a P.8 transposase plasmid (gifted by the Iain Cheeseman Lab). 24 hours after transfection, a media change was performed. After an additional 24 hours, cells were placed under G418 drug selection for 10 days, and then insertion was confirmed by microscopy.

Clones were first screened for loss of protein by immunoblotting. Genomic DNA was then isolated from validated clones, and the targeted locus was amplified by PCR. A list of primers is available in Supplementary Table 3. To confirm editing of all alleles, gel-purified PCR products were cloned using the TOPO TA Cloning Kit for Sequencing (Invitrogen; 450030), and individual colonies were selected for Sanger sequencing (Genewiz; **Supplementary Table 1**). Additionally, knockouts were validated in subsequently obtained WGS by manual inspection of read alignments using Integrative Genomics Viewer^41^.

APOBEC3C, APOBEC3F, and APOBEC3H knockdowns were established by generating mixed bulk knockout populations. Cells were transfected as described above for APOBEC3A and APOBEC3B deletions using 10 µg of the pU6-sgRNA_CBh-Cas9-T2A-mCherry plasmid encoding guides targeting the respective APOBEC3 paralogs (**Supplementary Table 2**). Twenty-four hours post-transfection, mCherry-positive cells were bulk sorted by FACS using the FACSAria system (BD Biosciences). RT-qPCR validated knockdown in bulk cultures.

### RNA isolation and reverse transcription quantitative PCR

RNA was isolated using the RNeasy Mini Kit (Qiagen; 74136). cDNA was synthesized from extracted RNA using Super-Script IV First-Strand Synthesis System (Invitrogen; 18091050). cDNA synthesis reactions were performed using 2 μl of 50 ng/μl random hexamers, 2 μl of 10 mM dNTPs, 4 μg RNA, and DEPC-treated water to a volume of 26 μl. A list of the primer sequences is provided in Supplementary Table 4.

### Immunoblotting

Cells were lysed in RIPA buffer (150 mM NaCl, 50 mM Tris-HCl pH 8.0, 1% NP-40, 0.5% sodium deoxycholate, 0.1% SDS, Pierce Protease Inhibitor Tablet, EDTA free) or sample buffer (62.5 mM Tris-HCl pH 6.8, 0.5 M β-mercaptoethanol, 2% SDS, 10% glycerol, 0.01% bromophenol blue). Quantification of RIPA extracts was performed using the Thermo Fisher Scientific Pierce BCA Protein Assay kit. Protein transfer was performed by wet transfer using 1x Towbin buffer (25 mM Tris, 192 mM glycine, 0.01% SDS, 20% methanol) and nitrocellulose membrane. Blocking was performed in 5% milk in 1x TBST (19 mM Tris, 137 mM NaCl, 2.7 mM KCl, and 0.1% Tween-20) for one hour at room temperature. The following antibodies were diluted in 1% milk in 1x TBST: anti-APOBEC3A/B/G (04A04; Millipore MABF3373; 1:1,000)^4^, anti-APOBEC3A (01D05; Millipore MABF3363; 1:1,000)^4^, anti-APOBEC3B (Abcam; ab184990; 1:500), anti-tubulin (Abcam; ab7291), anti-vinculin (Abcam; ab129002), anti-β-actin (Abcam, ab8227; 1:1,000); anti-mouse IgG HRP (Thermo Fisher Scientific; 31432; 1:10,000), and anti-rabbit IgG HRP (SouthernBiotech; 6441-05; 1:10,000).

### RNA-editing assay

*DDOST* 558C>U RNA-editing assays were performed as described previously^4,12^ with assistance from the MSKCC Integrated Genomics Operation. Total RNA was extracted using the RNeasy Mini kit (Qiagen; 74136) according to the manufacturer’s instructions. After extraction, cDNA was generated using the High Capacity cDNA Reverse Transcription Kit (Thermo Fisher Scientific; 4368814). cDNA (20 ng) along with primers purchased from Bio-Rad (10031279 and 10031276) for the target *DDOST558C>U* amplification was mixed with PCR reaction mix in a total volume of 25 µl. Then, 20 µL of the reaction was mixed with 70 µL of Droplet Generation Oil for Probes (Bio-Rad) and loaded into a DG8 cartridge (Bio-Rad). A QX200 Droplet Generator (Bio-Rad) was used to generate the droplets, which were transferred to a 96-well plate, and the following PCR reaction was then run: 5 min at 95 °C; 40 cycles of 94 °C for 30s and 53 °C for 1 min; and finally 98 °C for 10 min. The QX200 Droplet Reader (Bio-Rad) was used to analyze droplets for fluorescence measurements of the fluorescein amidite (FAM) and hexachlorofluorescein (HEX) probes. The data were analyzed using QuantaSoft (Bio-Rad), and gating was performed based on positive and negative DNA oligonucleotide controls.

### *in vitro* DNA deaminase activity assay

Cells (∼10^6^) were pelleted and lysed in deamination lysis buffer (25 mM HEPES (pH 8.0), 10% glycerol, 150 mM NaCl, 0.5% Triton X-100, 1 mM EDTA, Pierce™ Protease InhibitorCocktail). Lysates were sonicated using a Bioruptor 300 (Diagenode) with 15 cycles (30 s on, 30 s off) at 4°C or sheared through a 28 1/2-gauge syringe. Lysates were then cleared by centrifugation (e.g., 13,000 x g for 20 min at 4°C). Quantification of cleared lysates was performed using the Thermo Fisher Scientific Pierce BCA Protein Assay kit (Invitrogen; 23225). A total of 50 µg of protein was used for each time-course reaction.

Reactions were assembled in a final volume of 50 uL containing 50 µg lysate, 1x deaminase buffer (225 mM, Tris-HCl, pH 7.5, 20 mM KCl, 25 mM NaCl, 0.05% Triton X-100, 10 mM DTT), [0.1 µM] of the indicated 800-IRDye probe (TCA probe: 5 ′ IRD800/ T*T*T*GTAGATGTAGATGTAGTT**TCA**GTAGTAGAGTATGTAGT*A*T*T*T; or ACA probe: 5′IRD800/T*T*T*GTAGATGTAGATGTAGTT**ACA**GTAGTAGAGTATGTAGT*A*T*T*T), [1.5 units] of UNG (Uracil-DNA Glycosylase; NEB; M0372L), and ±0.5 µl RNase A (20 mg/ml; Thermofisher; 12091021). Reactions were incubated at 37 °C. At each time point, an aliquot was removed and treated with [2 µl of 2 M] NaOH, followed by incubation at 95 °C for 12 min to cleave abasic sites. Reactions were then neutralized with [2 µl of X 2 M] HCl.

Reactions were terminated by adding an equal volume of Novex™ TBE-Urea Sample Buffer (2X) (Thermofisher; LC6876) and separated on a 15% TBE-Urea gel (Thermofisher; EC68852BOX) in 1x TBE buffer for 120 min at 160 V. Gels were imaged using the Odyssey Infrared Imaging System (Li-COR) and quantified using ImageJ.

### Microscopy

For immunofluorescence microscopy, PC9 APOBEC3F-GFP cells were seeded on coverslips at 1×10^5^ cells/mL 24 hours before fixation and simultaneously treated with 1ug/mL doxycycline. Cells were washed three times with PBS prior to fixation in 2% paraformaldehyde (in PBS) for 15 minutes. After another PBS wash coverslips were incubated in permeabilization buffer (20 mmol/L Tris-HCl pH8, 50 mmol/L NaCl, 3 mmol/L MgCl2, 300 mmol/L Sucrose, 0.5% Triton X-100) for 10 minutes and then washed once more with PBS. The coverslips were incubated in blocking buffer ((1 mg/mL BSA, 3% goat serum, 0.1% Triton X-100, 1 mmol/L ethylenediamine tetraacetic acid (EDTA) in PBS) for 1 hour and GFP primary antibody (Abcam; ab290) diluted in blocking buffer was applied for 2 hours. Following three washes with PBS-TX (PBS, 0.1%Triton X-100), coverslips were incubated in the secondary antibody (Goat anti-Rabbit IgG Alexa Fluor 488, Invitrogen, A11034, 1:1,000), diluted in blocking buffer for 1 hour, and then washed three times with PBS-TX. DNA was stained with Hoechst 33342 (Thermo Fisher Scientific, 1 mg/mL) for 15 minutes followed by three PBS washes. Coverslips were then mounted in ProLong Gold Antifade Mountant (Life Technologies; P36930) Images were acquired on a Nikon Eclipse Ti2-E equipped with a CSU-W1 spinning disk with Borealis microadapter, Perfect Focus 4, motorized turret and encoded stage, polycarbonate thermal box, 5-line laser launch [405 (100 mW), 445 (45 mW), 488 (100 mW), 561 (80 mW), 640 (75 mW)], PRIME 95B Monochrome Digital Camera and 100X1.45 NA objective. Images were further edited with ImageJ and Adobe illustrator.

### Statistical analysis

Mutational statistical analyses were performed using R 4.1.2 unless otherwise specified. Wilcoxon rank-sum tests were used to test for significance, and multiple test correction was performed using Bonferroni correction unless otherwise specified.

### DNA extraction and WGS

Genomic DNA was extracted from individual clones using the DNeasy Blood and Tissue Kit (Qiagen, 51106). dsDNA was quantified using the Qubit dsDNA BR Quantification Assay kit (Thermofisher, Q32850), and results were read on a Qubit 4Fluorometer (Thermofisher, Q33238). Extracted DNA from each clone was subjected to 150bp paired-end WGS on Illumina NovaSeq platforms (‘6000’ and ‘X’), achieving an average fold-coverage of approximately 30× per sample.

### WGS analysis

Reads containing sequencing adapters were identified and marked using Genome Analysis Toolkit (GATK) MarkIlluminaAdapters^42^. The marked reads were mapped to the human reference genome GRCh37^43^ using bwa-mem version 0.7.17-r1188 ^44^ and converted to binary alignment map format using Samtools version 1.18^45^. GATK MergeBamAlignment was used to merge the BWA output with the output of MarkIlluminaAdapters. GATK MarkDuplicatesSpark was used to mark duplicate reads. Alignment files for each sample were merged and converted into compressed reference-oriented alignment map (CRAM) files using GATK MergeSam and Samtools for storage and downstream analysis. Version 4.4.0.0 was used across all cited GATK tools.

### Mutation calling

Identifying *de novo* SBS and indel mutations in cell lines can be complicated by low-frequency pre-existing somatic variants in the parental lineage that arose prior to the examined propagation times and may evade consistent classification by standard variant callers. To address this, we employed a two-pronged variant calling strategy to identify and exclude germline and pre-existing somatic mutations. For each daughter clone, mutations were called using GATK Mutect2 (version 4.4.0.0) with the parental clone specified as the matched normal, using default parameters^46^(*default* run). Because this mode does not report variants classified as likely germline, we additionally ran Mutect2 with --force-call-filtered-alleles and --genotype-germline-sites (*comprehensive* run) to recover these sites. Calls from both runs were merged to generate a unified mutation set per daughter clone. The merged mutation sets obtained from the default and comprehensive Mutect2 runs were treated as an initial call set and subjected to additional post hoc filtering to ensure retention of *de novo* variants only. First, SBS and indel mutations whose genomic locus was covered by <15 reads in the matched parental clone were excluded, as insufficient depth precludes confident exclusion of pre-existing variants. Second, mutations supported by at least one read in more than 50% of parental clones from the same cell line were removed, under the assumption that such variants were pre-existing but stochastically missed in the matched parent due to sampling variability. Third, mutations observed in at least one read in daughter clones derived from different parental lineages were excluded; these sites were frequently located in repetitive regions and exhibited mutational profiles inconsistent with bona fide private mutations. The remaining variants were retained as high-confidence *de novo* mutations and used for all downstream analyses. Mutations in parent clones identified using Mutect2 version 4.4.0.0^46^, calling against the reference genome.

### Cell line validation and sample swap detection

The identity of all sequenced cell lines was validated by SNP genotyping. For each sample, we genotyped a panel of 93 SNP sites previously established for fingerprinting hundreds of human cell lines in the COSMIC Cell Line Project (https://cancer.sanger.ac.uk/cancergenome/assets/cell_lines/QC.xlsx) using BCFtools ^45^.

To identify potential sample swaps—instances in which a daughter clone was derived from a different parental clone than annotated—we first identified mutations private to individual parental clones using GATK Mutect2 without a matched normal. Variants detected in more than one parental clone were removed, as were variants with a variant allele frequency (VAF) <0.25, yielding a catalog of private mutations for each parental clone. Daughter clones were then genotyped at these private parental variant sites to assess relatedness to each parental clone. We determined a parent-daughter pair to be related if the daughter clone harbored reads supporting >50% of private parental mutations. Identified swaps were corrected by reassigning the annotated parent for the affected daughter clones where possible. Daughter clones for which no parental clone could be confidently assigned were excluded from downstream analyses.

### Analysis of *de novo* acquired mutations and mutational signatures in cell lines lineages

SBS mutations from daughter clones were converted into mutational catalogs summarizing the number of mutations in particular mutational contexts using SigProfilerMatrixGenerator v1.2.31^45,47^. We generated 288-context catalogs, which classify mutations by substitution type, the mutated pyrimidine base, the immediate 5′ and 3′ flanking nucleotides, and transcriptional strand. *De novo* mutational signature extraction was performed using SigProfilerExtractor version 1.1.24^48^. Extracted signatures were subsequently assigned to canonical COSMIC mutational signatures^3^ using SigProfilerAssignment version 0.1.6, enabling quantification of each canonical signature in each daughter clone ^49^.

Extended SBS sequence context enrichment was calculated as described previously^9^. Briefly, counts of YTCA and RTCA context mutations were obtained from SigProfilerMatrixGenerator outputs. Genome-wide frequencies of YTCA and RTCA contexts were quantified using Jellyfish^50^ and used to normalize mutation counts. Enrichment scores for YTCA and RTCA mutations were calculated for each sample as described^9^. Clusters of SBS mutations were identified using SigProfilerClusters^51^ version 1.1.2.

We generated catalogs of indel mutations using the 93-channel indel classification paradigm^21^ with indelsig.tools.lib, commit 4a293b0^21^. Mutational signatures were extracted from these catalogs with SigProfilerExtractor^48^ version 1.1.24 and were decomposed into the reference tumor signatures^21^ using non-negative least squares implemented through the glmnet package in R^52^, using an alpha value of 0.1.

## Data Availability

Raw sequencing data have been deposited in the NCBI Sequence Read Archive under the BioProject ID PRJNA1366483.

## Acknowledgements

We thank members of the Maciejowski and Petljak labs for work on this manuscript. Work in J.M.’s laboratory is supported by the NCI (R37CA261183; R01CA270102; R01CA304441; P30CA008748), the Pershing Square Sohn Cancer Research Alliance, the Allen and Sandra Gerry Metastasis and Tumor Ecosystems Center, the Frank A. Howard Scholars Program, the Mary Kay Ash Foundation and the Experimental Therapeutics Centers at MSKCC. J.S. is supported by the NCI (F31CA290850). Work in M.P.’s laboratory for this study was supported by R01CA270102.

## Contributions

Conceptualization: J.S., M.P., and J.M.; Methodology: J.S., M.P. and J.M.; Investigation: J.S. and L.C.; Formal Analysis: J.S. and L.C.; Writing - Original Draft: J.S. and J.M.; Writing - Review and Editing - all authors; Visualization: J.S. and J.M.; Funding Acquisition: J.S., M.P. and J.M; Supervision: M.P. and J.M.

