## Supplementary Information for "A context dependent hierarchy of APOBEC3A and APOBEC3B mutators in lung adenocarcinoma"

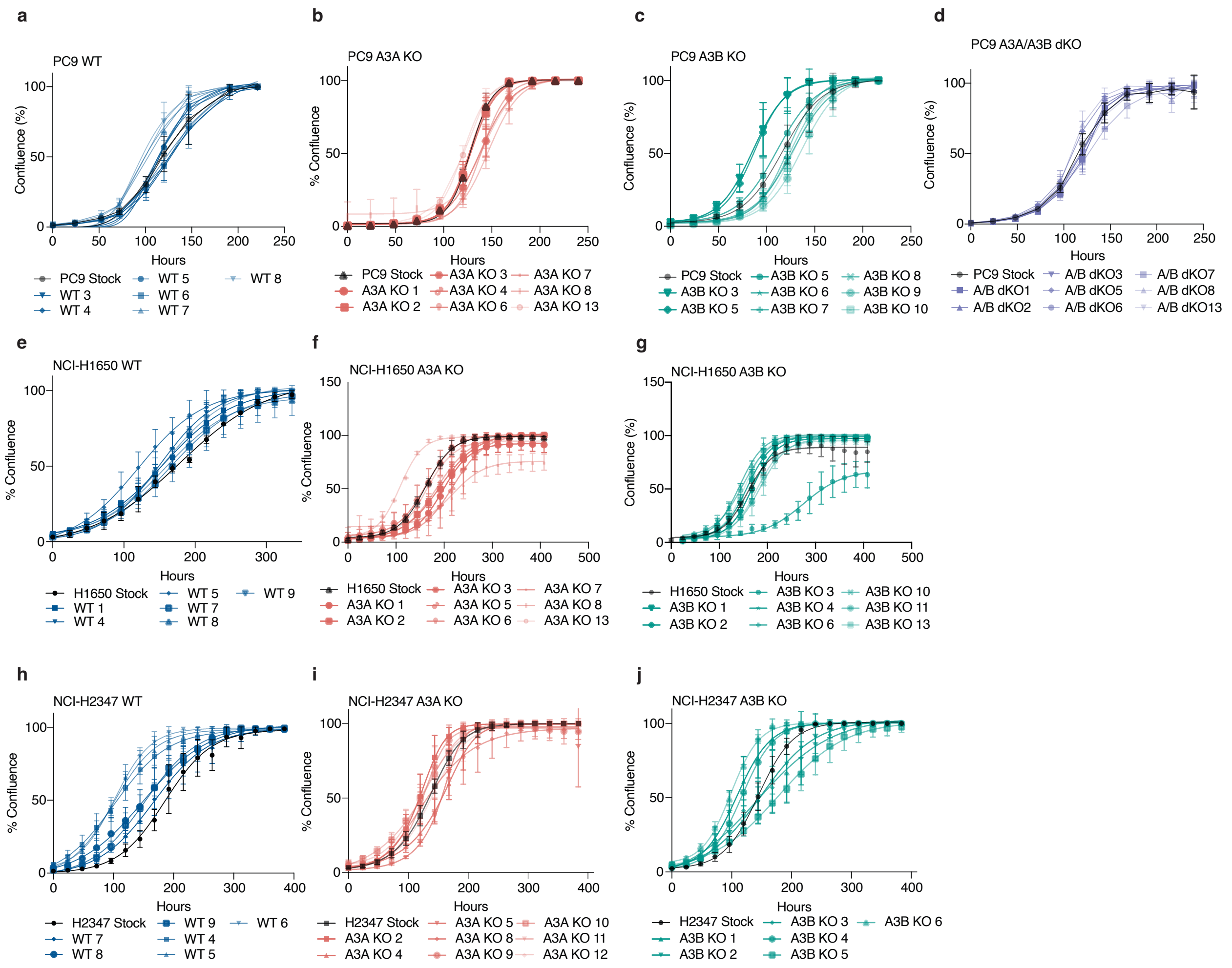


**Supplementary Figure 1. Proliferation rates of single-cell derived LUAD lineages.**
Cell confluency, measured over time, is shown for the parental population (black line) and the individual parental clones (colored lines) used for long-term propagation. Panels are grouped by cell line and genotype: (a-d) PC9 clones: (a) WT, (b) *APOBEC3A* KO (A3A KO), (c) *APOBEC3B* KO (A3B KO), (d) *APOBEC3A/APOBEC3B* dKO (A3A/A3B dKO). (e-g) NCI-H1650 clones: (e) WT, (f) *APOBEC3A* KO, (g) *APOBEC3B* KO. (h-j) NCI-H2347 clones: (h) WT, (i) *APOBEC3A* KO, (j) *APOBEC3B* KO. Each growth curve represents a single clone, plotted as mean ± s.d. of $n = 3$ technical replicates.

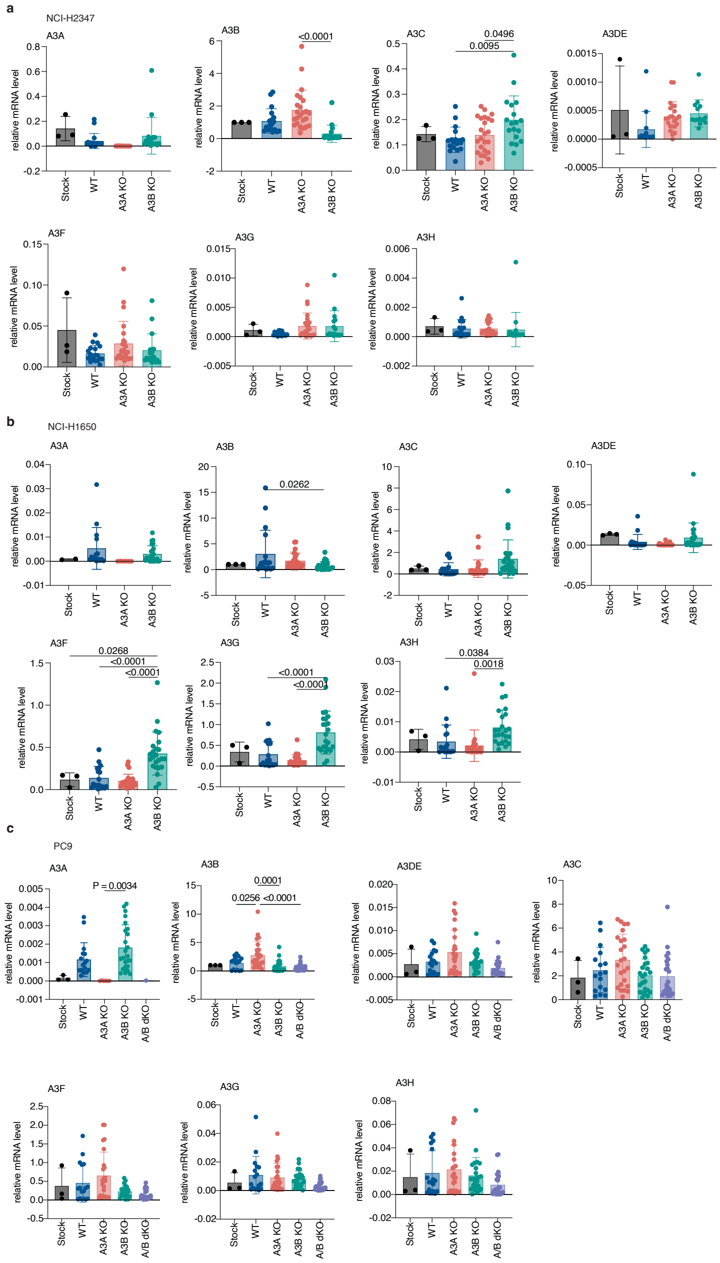


**Supplementary Figure 2. Knockout of *APOBEC3A* or *APOBEC3B* does not induce compensatory expression of other APOBEC3 family members.**

qPCR analysis of relative mRNA levels for all *APOBEC3* family members in (a) NCI-H2347, (b) NCI-H1650, and (c) PC9. Expression levels are shown for the parental population, panels of wild-type (WT) clones, and the *APOBEC3A* KO (A3A KO), *APOBEC3B* KO (A3B KO), and *APOBEC3A/APOBEC3B* dKO (A3A/A3B dKO) clones used in this study. The analysis confirms loss of the targeted transcript without evidence of systematic compensatory upregulation of other *APOBEC3* family members. Data are shown as mean ± s.d., with each dot representing an independent biological replicate; 6-8 clones per genotype are grouped. *P* values were calculated using an ordinary one-way ANOVA with Tukey's multiple-comparisons test.

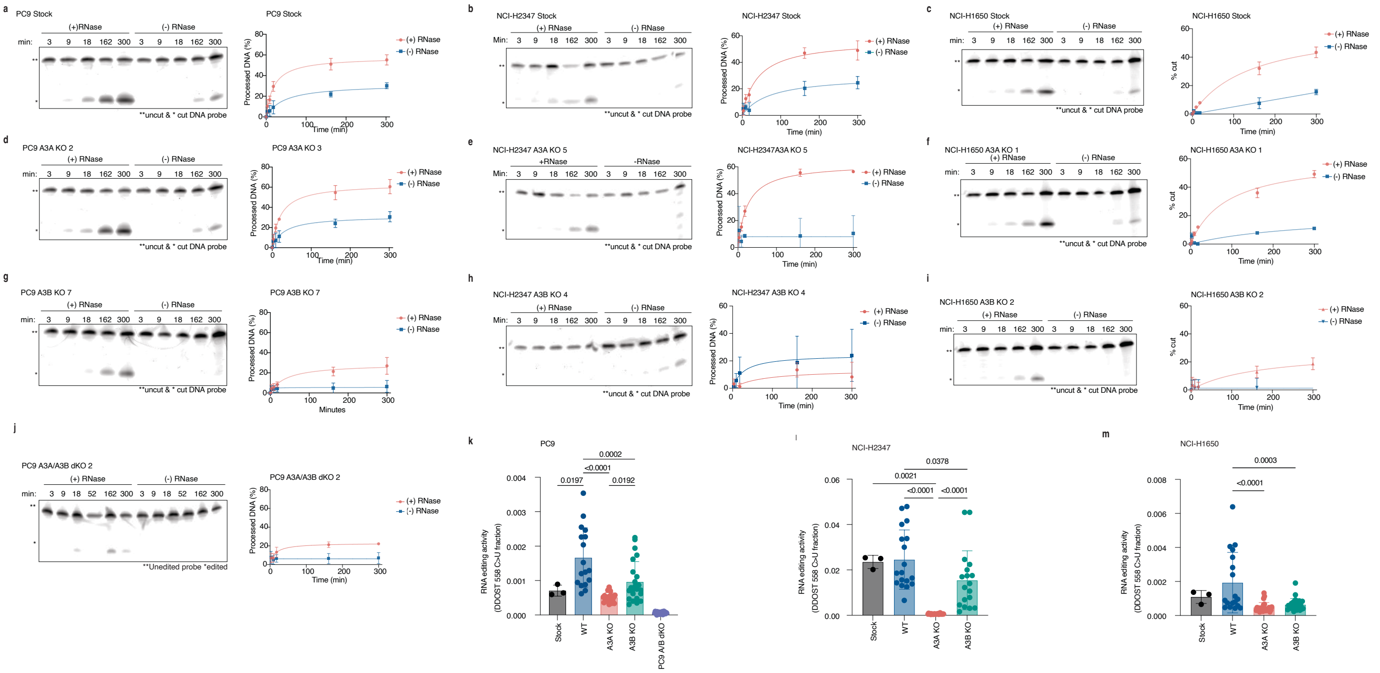


**Supplementary Figure 3. Functional characterization of APOBEC3 knockout clones.**
(a-j) *In vitro* DNA deaminase activity in whole-cell lysates from parental and the indicated KO clones. Activity was measured on a linear oligonucleotide substrate over a 300-minute time course, either with (+RNase A, red line) or without (-RNase A, blue line) RNase A treatment. Representative gels are shown to the left of each plot. Data are presented as mean ± s.d. from three independent biological replicates. (k-m) Quantification of RNA editing activity at the *DDOST* C558 hotspot in parental, WT, and KO clones for (k) PC9, (l) NCI-H2347, and (m) NCI-H1650. Data are shown as mean ± s.d., with each dot representing an independent biological replicate; 6-8 clones per genotype are grouped. *P* values were calculated using an ordinary one-way

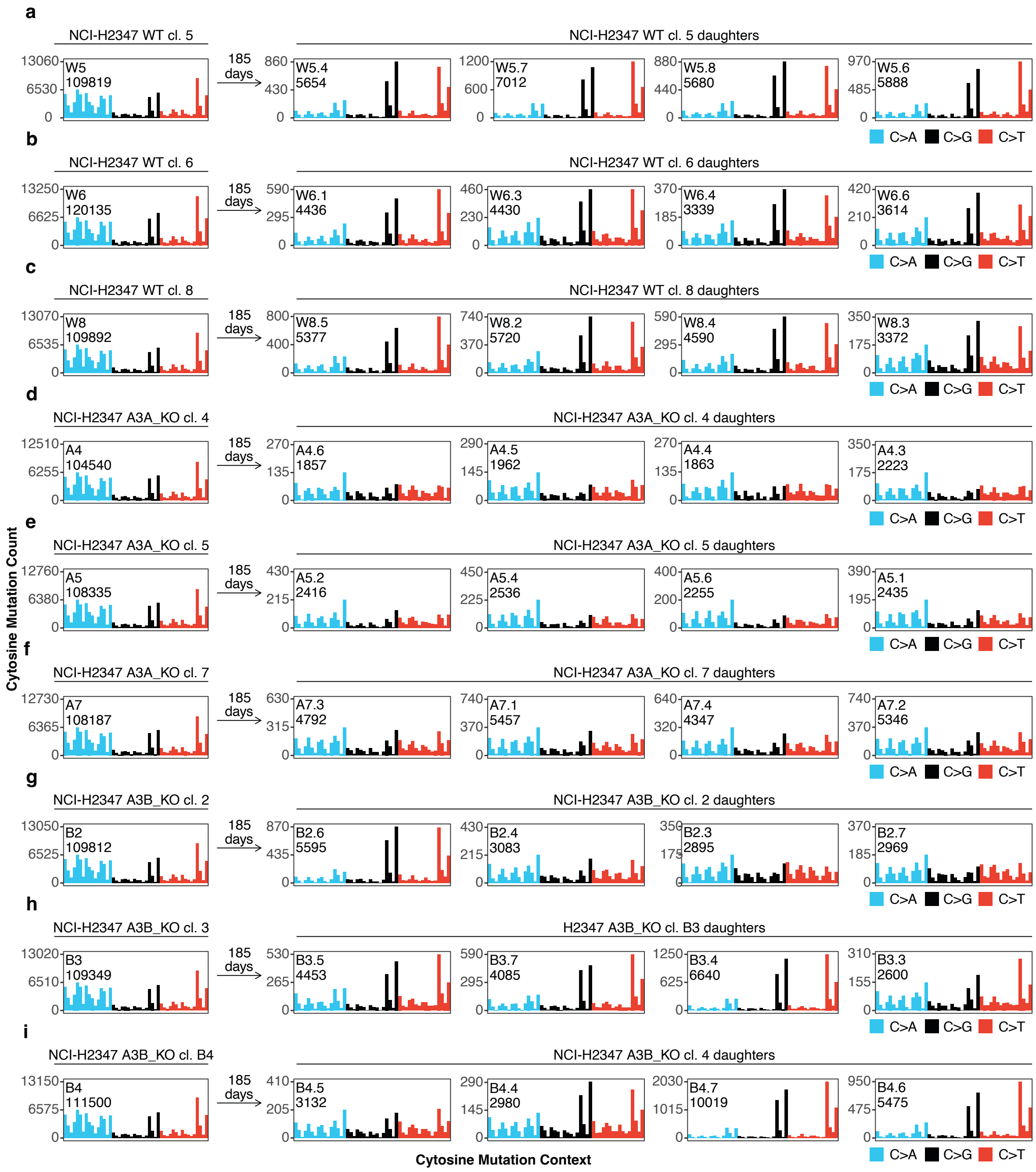


**Supplementary Figure 4. Patterns of *de novo* acquired SBS in NCI-H2347 cell lineages.**
Profiles of *de novo* SBS mutations acquired in the indicated daughter clone lineages and of mutations identified in their corresponding parental clone. Each plot is displayed as the number of mutations (y-axis) attributed to 48 cytosine SBS classes (x-axis), which are defined by the 3 color-coded substitution classes and sequence context immediately 3′ and 5′ to the mutated base. The plots are grouped according to the parental clone from which each set of single-cell derived daughter clone lineages was derived: a–c, wild-type (WT) lineages; d–f, *APOBEC3A* knockout (A3A KO) lineages; and g–i, *APOBEC3B* knockout (A3B KO) lineages. *De novo* mutations identified in daughter lineages from NCI-H2347 were acquired during 185 days of culture.

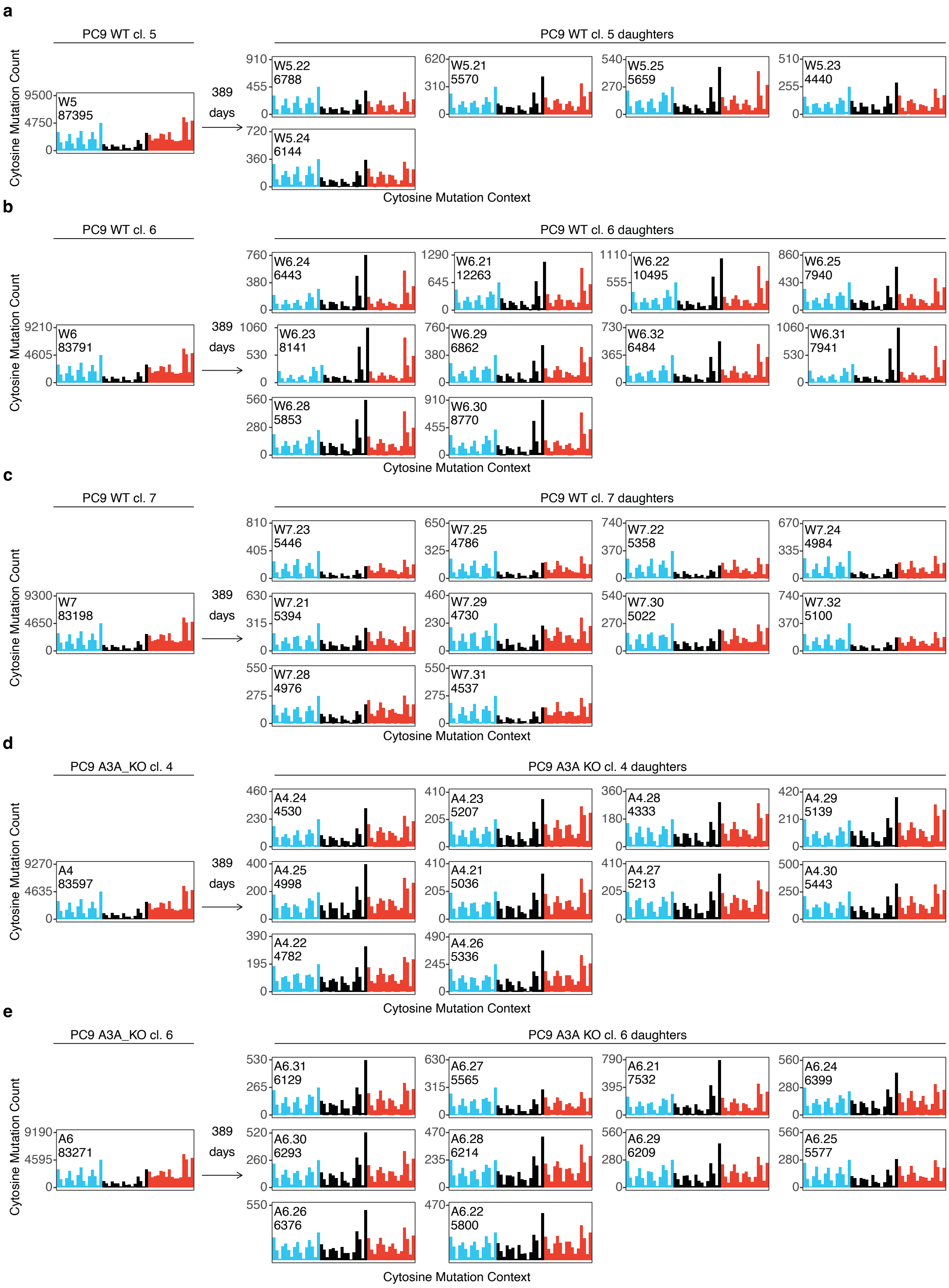
a
PC9 WT cl. 5
Cytosine Mutation Count
W5
87395
9500
4750
0
389
days
PC9 WT cl. 5 daughters
W5.22
6788
W5.21
5570
W5.25
5659
W5.23
4440
W5.24
6144
Cytosine Mutation Context
b
PC9 WT cl. 6
W6
83791
PC9 WT cl. 6 daughters
W6.24
6443
W6.21
12263
W6.22
10495
W6.25
7940
W6.23
8141
W6.29
6862
W6.32
6484
W6.31
7941
W6.28
5853
W6.30
8770
c
PC9 WT cl. 7
W7
83198
PC9 WT cl. 7 daughters
W7.23
5446
W7.25
4786
W7.22
5358
W7.24
4984
W7.21
5394
W7.29
4730
W7.30
5022
W7.32
5100
W7.28
4976
W7.31
4537
d
PC9 A3A_KO cl. 4
A4
83597
PC9 A3A KO cl. 4 daughters
A4.24
4530
A4.23
5207
A4.28
4333
A4.29
5139
A4.25
4998
A4.21
5036
A4.27
5213
A4.30
5443
A4.22
4782
A4.26
5336
e
PC9 A3A_KO cl. 6
A6
83271
PC9 A3A KO cl. 6 daughters
A6.31
6129
A6.27
5565
A6.21
7532
A6.24
6399
A6.30
6293
A6.28
6214
A6.29
6209
A6.25
5577
A6.26
6376
A6.22
5800

**Supplementary Figure 5. Patterns of *de novo* acquired SBS in PC9 cell lineages.**

Profiles of *de novo* SBS mutations acquired in the indicated daughter clone lineages and of mutations identified in their corresponding parental clone. Each plot is displayed as the number of mutations (y-axis) attributed to 48 cytosine SBS classes (x-axis), which are defined by the 3 color-coded substitution classes and sequence context immediately 3′ and 5′ to the mutated base. The plots are grouped according to the parental clone from which each set of single-cell derived daughter clone lineages was derived: a–c, wild-type (WT) lineages; d, e, *APOBEC3A* knockout (A3A KO) lineages; f–h, *APOBEC3B* knockout (A3B KO) lineages; and i–k, *APOBEC3A/APOBEC3B* double knockout (A/B dKO) lineages. *De novo* mutations identified in daughter lineages from PC9 were acquired during 389 days of culture.

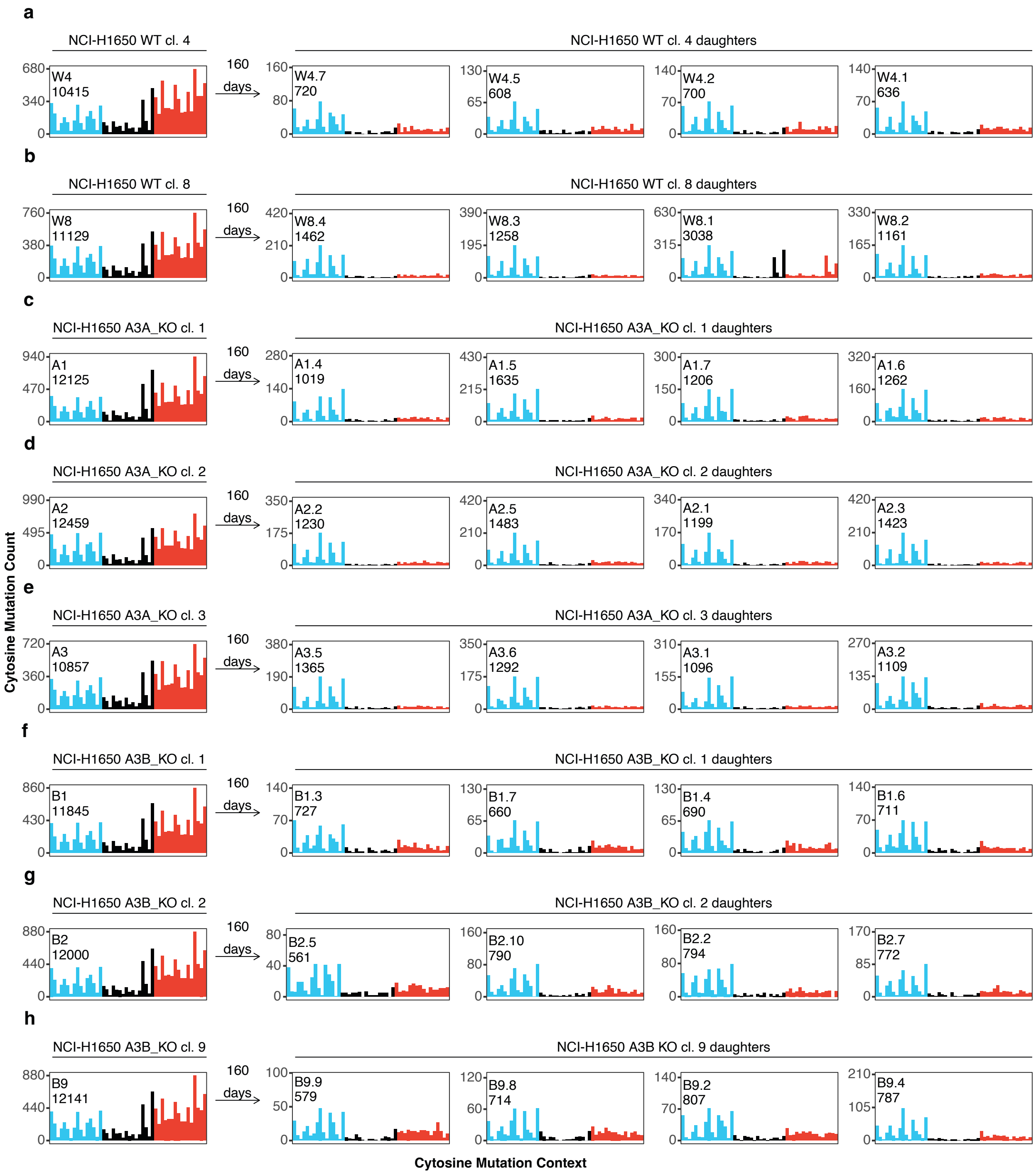


**Supplementary Figure 6. Patterns of *de novo* acquired SBS in NCI-H1650 cell lineages.**
Profiles of *de novo* SBS mutations acquired in the indicated daughter clone lineages and of mutations identified in their corresponding parental clone. Each plot is displayed as the number of mutations (y-axis) attributed to 48 cytosine SBS classes (x-axis), which are defined by the 3 color-coded substitution classes and sequence context immediately 3′ and 5′ to the mutated base. The plots are grouped according to the parental clone from which each set of single-cell derived daughter clone lineages was derived: a, b, wild-type (WT) lineages; c–e, *APOBEC3A* knockout (A3A KO) lineages; and f–h, *APOBEC3B* knockout (A3B KO) lineages. *De novo* mutations identified in daughter lineages from NCI-H1650 were acquired during 160 days of culture.

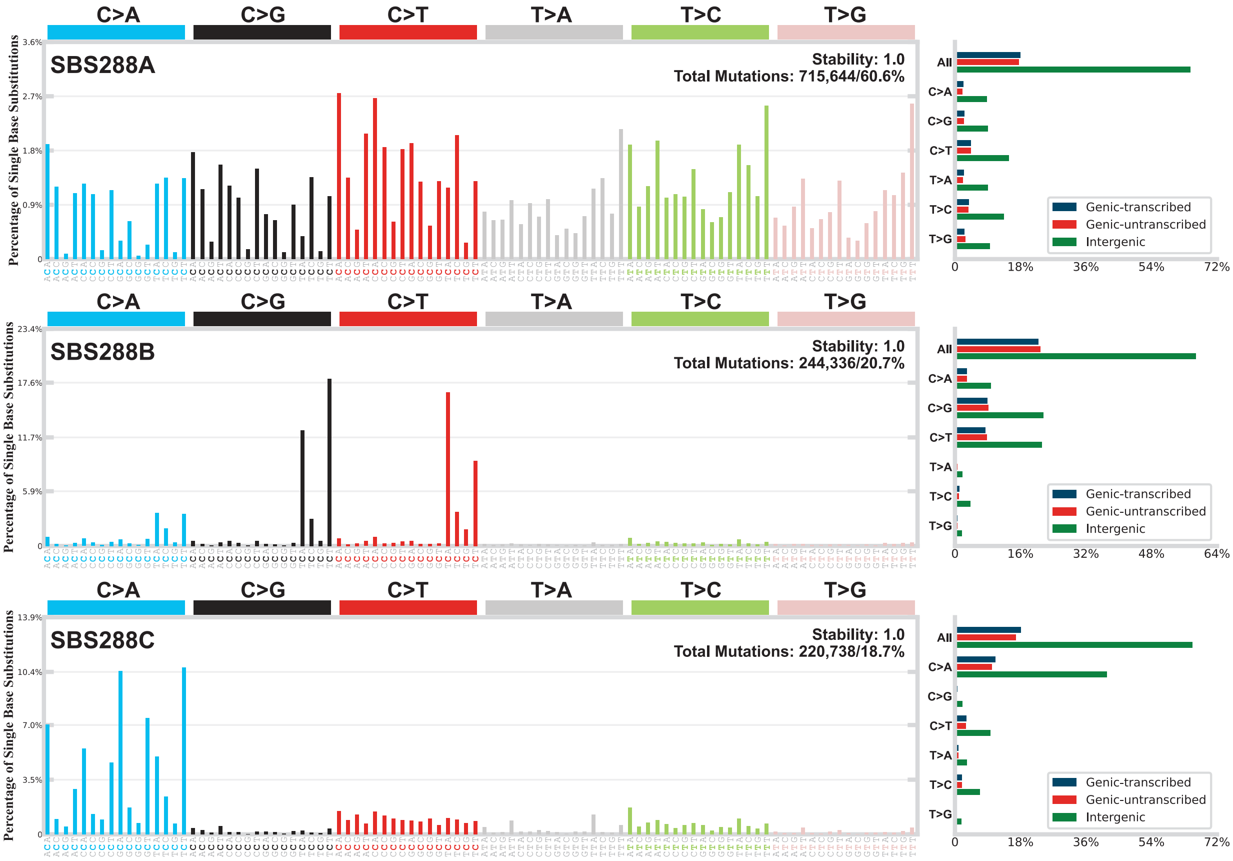


**Supplementary Figure 7. SBS signatures identified among mutations acquired in examined LUAD lineages.**

SBS signatures extracted from mutational catalogs of *de novo* mutations identified across 175 LUAD daughter clones from PC9, NCI-H2347 and NCI-H1650 cell lines using SigProfilerExtractor. Each signature is displayed in standardized 288-channel mutational contexts.

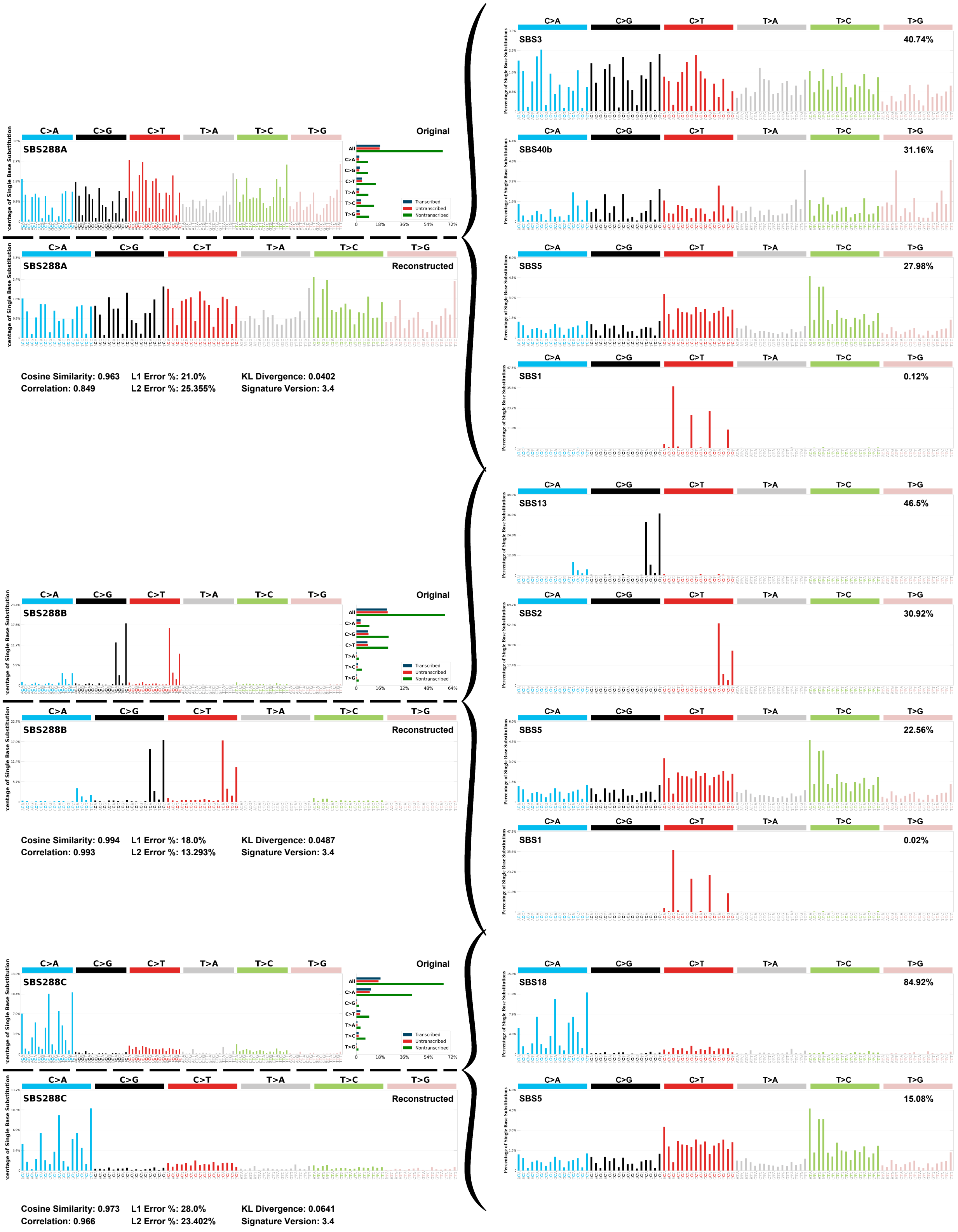


**Supplementary Figure 8. Decomposition of *de novo* extracted SBS signatures into COSMIC tumor census signatures**

SigProfilerAssignment was used to fit census SBS signatures to the signatures extracted from *de novo* mutations identified in daughter clones. Decomposition of *de novo* SBS signature SBS288B revealed contributions from the APOBEC3-associated signatures SBS2/13.

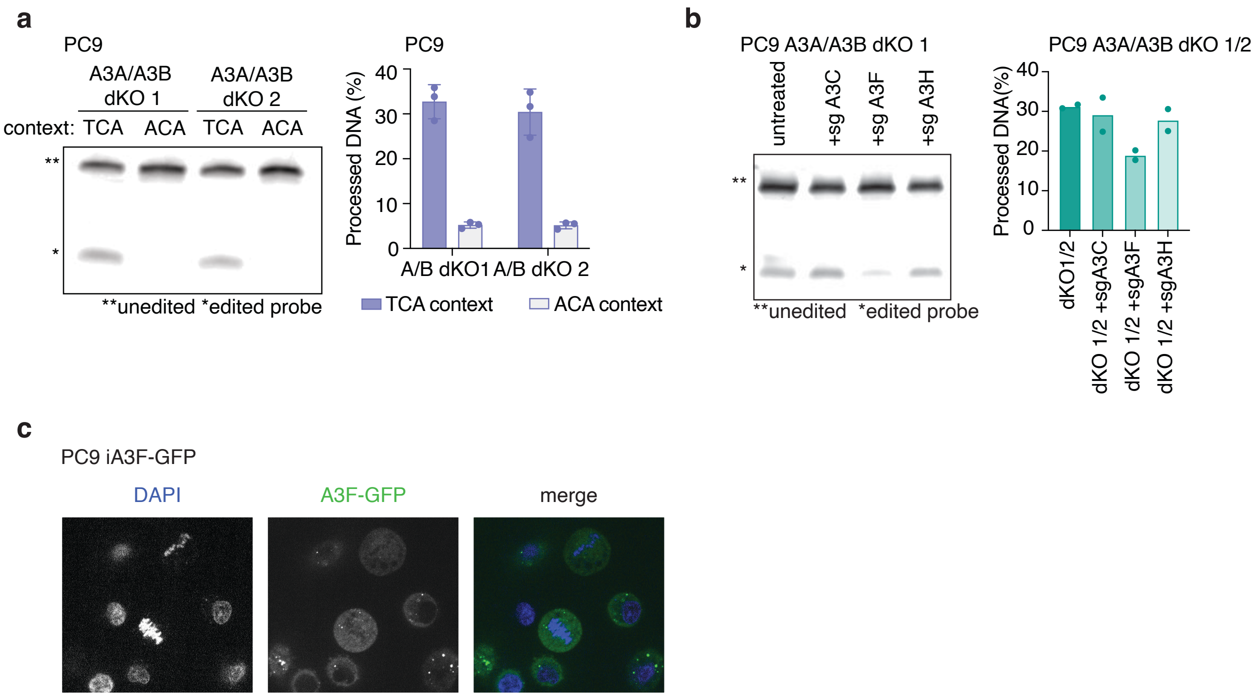


**Supplementary Figure 9. Residual deaminase activity in PC9 *APOBEC3A/APOBEC3B* dKO clones is driven by cytosolic APOBEC3F.**

(a) *In vitro* DNA deaminase activity of lysates from two independent PC9 *APOBEC3A/APOBEC3B* dKO (*A3A*/*A3B* dKO) clones. Activity was measured on oligonucleotide substrates containing either the APOBEC3-preferred TCA motif or a disfavored ACA motif. The representative gel (left) and quantification (right) are shown. (b) *In vitro* DNA deaminase activity in two PC9 *APOBEC3A/APOBEC3B* dKO (*A3A*/*A3B* dKO) clones following CRISPR-mediated knockdown of *APOBEC3C (A3C)*, *APOBEC3F (A3F)*, or *APOBEC3H (A3H)*. The representative gel (left) and quantification (right) are shown. (c) Immunofluorescence microscopy of PC9 cells expressing an APOBEC3F-GFP fusion protein (green). Nuclei are counterstained with DAPI (blue). The merged image shows APOBEC3F-GFP is excluded from the nucleus. For all deaminase assays (a, b), gels are representative of two independent biological replicates. Bar plots show mean ± s.d. for $n$ = 2 biological replicates (a), or n=1 (b).

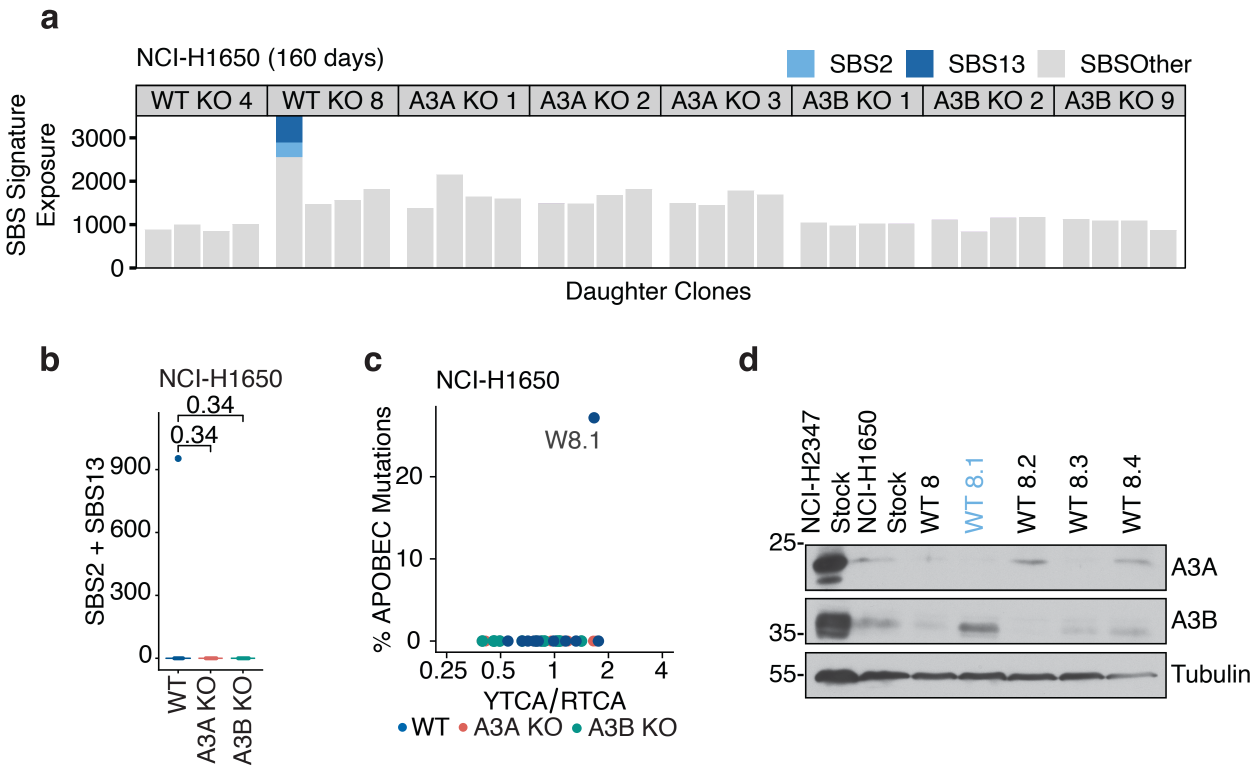


**Supplementary Figure 10. NCI-H1650 exhibits rare APOBEC3-mediated mutagenic episodes.**
(a) *De novo* SBS signature exposure in individual NCI-H1650 daughter clones (WT, A3A KO, A3B KO) following 160 days propagation. Signatures are color-coded: SBS2 (light blue), SBS13 (dark blue), and other SBS (grey). (b) Analysis of *de novo* SBS2/13 mutational burdens acquired in NCI-H1650 clones across annotated genotypes reveals that only a single wild-type (WT) lineage exhibited detectable APOBEC3-associated mutagenic activity, as evidenced by SBS2/13 mutations. (c) Enrichment of *de novo* APOBEC3-associated mutations from NCI-H1650 daughter clones in APOBEC3B-preferred RTCA vs APOBEC3A-preferred YTCA sequence contexts. Each point represents one clone, colored by genotype. (d) Immunoblot analysis of APOBEC3A (A3A) and APOBEC3B (A3B) protein levels in NCI-H1650 WT daughter clones. Lysates from NCI-H2347 (APOBEC3A-high) and NCI-H1650 (parental) are included as controls. Tubulin serves as a loading control. Box plots in (b) show the median (center line), 25th and 75th percentiles (box limits), and whiskers extending to 1.5 x interquartile range. Each dot represents the combined SBS2 and SBS13 mutation burdens in each individual daughter clone. *P*-values shown were calculated using a two-tailed Mann-Whitney U test comparing SBS2+SBS13 burdens in each knockout genotype to the wild-type (WT) of the same cell line and time point.

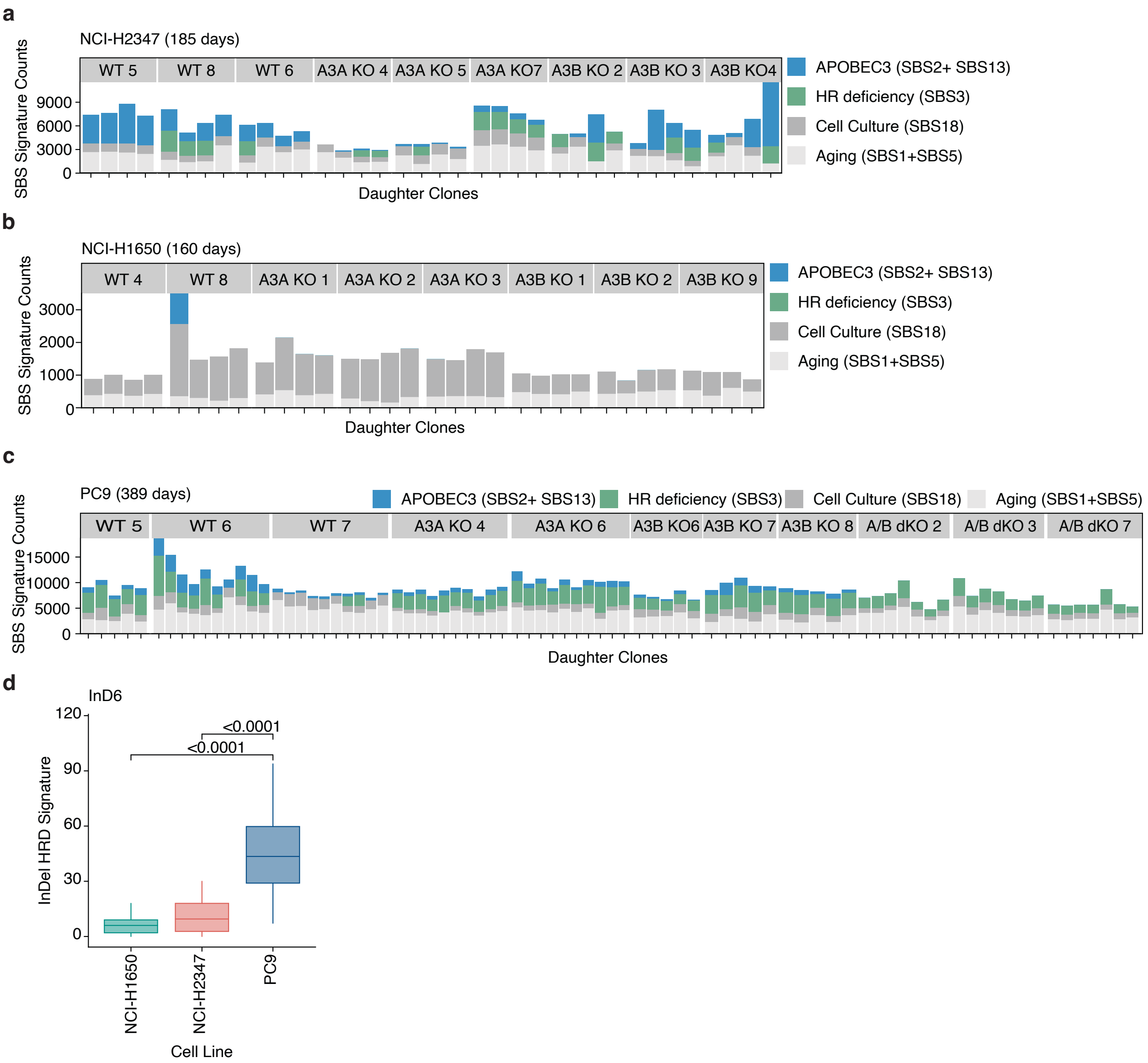


**Supplementary Figure 11. Quantification of mutational signature burdens across examined LUAD lineages.** Stacked bar plots showing the total number of de novo single-base substitution (SBS) mutations (y-axis) attributed to the indicated mutational signatures across individual daughter clones (x-axis). Signatures are color-coded as follows: APOBEC3 (SBS2+13, blue), homologous recombination (HR) deficiency (SBS3, green), cell culture–associated (SBS18, dark grey), and aging-associated (SBS1+5, light grey). (a) NCI-H2347 daughter clones, (b) NCI-H1650 daughter clones (same data as in Supplementary Figure 10a), and (c) PC9 daughter clones. (d) Quantification of *de novo* InD6a mutation accumulation in WT daughter clones from PC9, NCI-H1650, and NCI-H2347. Box plots show the median (center line), 25th and 75th percentiles (box limits), and whiskers extending to the most extreme data points. Each dot represents an individual daughter clone. P-values shown were calculated using a two-tailed Mann-Whitney U test comparing LUAD cell lines.

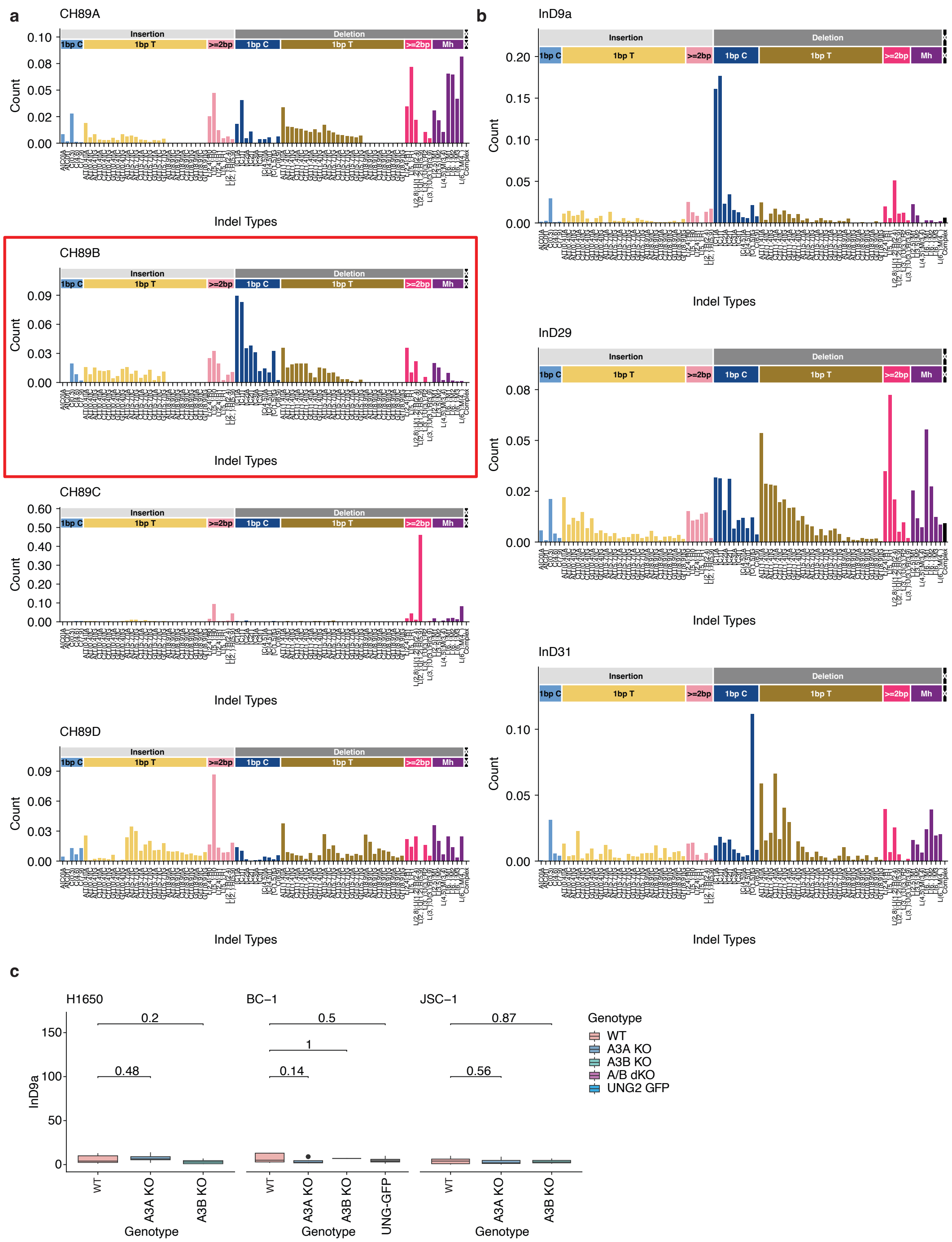


**Supplementary Figure 12. Indel signatures identified among mutations acquired in examined LUAD lineages.**

(a) Mutational profiles of indel signatures *de novo* extracted and refitted from mutational catalogs of LUAD daughter clones using SigProfilerExtractor and non-negative least squares refitting. Bars represent probabilities (y-axis) of indel mutations in 89 sequence contexts (x-axis) as defined in Koh et al. De novo signature containing InD9a outlined in red. (b) Decomposition of CH89B using reference InDel mutational signatures. Reference mutational signatures (Koh et al, 2025) were fitted to each de novo signature using non-negative least squares. Representative refitting results from CH89B, including presumptive APOBEC3A signature InD9a are shown. (c) Quantification of *de novo* InD9a mutation accumulation in daughter clones from NCI-H1650 (left), BC-1 (center), and JSC-1 (right) cell lines. Box plots show the median (center line), 25th and 75th percentiles (box limits), and whiskers extending to 1.5 times x interquartile range. Dots indicate outlier clones with InD9a mutation burdens beyond the whisker range. P-values shown were calculated using a two-tailed Mann-Whitney U test comparing each genotype to its respective wild-type (WT) control.

| Cell Line | % colonies | Description |
|---|---|---|
| **PC9 A3A KO 1** | 100 | 72 bp deletion leading to premature stop codon at cut site (aa12-36 exon 3; bp38959547 to 38959618) |
| **PC9 A3A KO 2** | 100 | 72 bp deletion leading to premature stop codon at cut site (aa12-36 exon 3; bp38959547 to 38959618) |
| **PC9 A3A KO 3** | 100 | 72 bp deletion leading to premature stop codon at cut site (aa12-36 exon 3; bp38959547 to 38959618) |
| **PC9A3A KO 4** | 100 | 72 bp deletion leading to premature stop codon at cut site (aa12-36 exon 3; bp38959547 to 38959618) |
| **PC9 A3A KO 5** | 90% | 72 bp deletion leading to premature stop codon at cut site (aa12-36 exon 3; bp38959547 to 38959618) |
| | 10% | 74 bp deletion causing the rest of exon 3 and 4 to be scrambled and an early stop midway through exon 4 (START 38959545 END 38959618 LENGTH 74) |
| **PC9 A3A KO 6** | 62.50% | 72 bp deletion leading to premature stop codon at cut site (aa12-36 exon 3; bp38959547 to 38959618) |
| | 12.50% | 77 bp deletion causing the rest of exon 3 and 4 to be scrambled and an early stop midway through exon 4 (START 38959545 END 38959618 LENGTH 74) |
| | 12.50% | 73 bp deletion causing the rest of exon 3 and 4 to be scrambled and an early stop midway through exon 4(START 38959546 END 38959618 LENGTH 73bp) |
| | 12.50% | 71 bp deletion leading to premature stop codon at cut site (aa12-36 exon 3; bp38959547 to 38959617) |
| **PC9 A3A KO 7** | 90% | 72 bp deletion leading to premature stop codon at cut site (aa12-36 exon 3; bp38959547 to 38959618) |
| | 10% | 71 bp deletion leading to premature stop codon at cut site (aa12-36 exon 3; bp38959547 to 38959617) |
| **PC9 A3A KO 8** | 77.78% | 72 bp deletion leading to premature stop codon at cut site (aa12-36 exon 3; bp38959547 to 38959618) |
| | 11.11% | 71 bp deletion leading to premature stop codon at cut site (aa12-36 exon 3; bp38959547 to 38959617) |
| | 11.11% | 76 bp deletion leading to complete scramble of exon 3 and early stop codon at the beginning of exon 4 (START 38959542 END 38959618 LENGTH 77) |
| **PC9 A3A KO 9** | 80% | 72 bp deletion leading to premature stop codon at cut site (aa12-36 exon 3; bp38959547 to 38959618) |
| | 20% | 71 bp deletion leading to premature stop codon at cut site (aa12-36 exon 3; bp38959547 to 38959617) |
| **H1650 A3A KO 1** | 100% | 72 bp deletion leading to premature stop codon at cut site (aa12-36 exon 3; bp38959547 to 38959618) |
| **H1650 A3A KO 2** | 83.33% | 71 bp deletion leading to premature stop codon at cut site (aa12-36 exon 3; bp38959547 to 38959617) |
| | 16.67% | 72 bp deletion leading to premature stop codon at cut site (aa12-36 exon 3; bp38959547 to 38959618) |
| **H1650 A3A KO 3** | 100% | 72 bp deletion leading to premature stop codon at cut site (aa12-36 exon 3; bp38959547 to 38959618) |
| **H1650 A3A KO 5** | 100% | 72 bp deletion leading to premature stop codon at cut site (aa12-36 exon 3; bp38959547 to 38959618) |
| **H1650 A3A KO 6** | 100% | 162 bp deletion leading to premature stop codon at the beginning of exon 4 |
| **H1650 A3A KO 7** | 50% | 12 bp deletion right over guide 4 leading to premature stop codon in exon 3 |
| | 50% | 72 bp deletion leading to premature stop codon at cut site (aa12-36 exon 3; bp38959547 to 38959618) |
| **H1650 A3A KO 8** | 50% | 71 bp deletion leading to premature stop codon at cut site (aa12-36 exon 3; bp38959547 to 38959617) |
| | 50% | 72 bp deletion leading to premature stop codon at cut site (aa12-36 exon 3; bp38959547 to 38959618) |

| | | |
|---|---|---|
| **H1650 A3A KO 13** | 100% | 72 bp deletion leading to premature stop codon at cut site (aa12-36 exon 3; bp38959547 to 38959618) |
| **H2347 A3A KO 4** | 100% | 72 bp deletion leading to premature stop codon at cut site (aa12-36 exon 3; bp38959547 to 38959618) |
| **H2347 A3A KO 5** | 88.88% | 72 bp deletion leading to premature stop codon at cut site (aa12-36 exon 3; bp38959547 to 38959618) |
| | 11.12% | two small deletions one on guide 3 and one on guide 4 (first deletion gets rid of splice site second one causes scramble and early stop codon in exon 4) |
| **H2347 A3A KO 7** | 11.12% | single base deletion: deletion of the middle A in E36 causing a frame shift and early stop codon in exon 4 |
| | 88.88% | 72 bp deletion leading to premature stop codon at cut site (aa12-36 exon 3; bp38959547 to 38959618) |
| **H2347 A3A KO 8** | 100% | 72 bp deletion leading to premature stop codon at cut site (aa12-36 exon 3; bp38959547 to 38959618) |
| **H2347 A3A KO 10** | 20% | two small deletions one on guide 3 and one on guide 4 (first deletion gets rid of splice site second one causes scramble and early stop codon in exon 4) |
| | 50% | 71 bp deletion leading to premature stop codon at cut site (aa12-36 exon 3; bp38959547 to 38959617) |
| | 30% | 72 bp deletion leading to premature stop codon at cut site (aa12-36 exon 3; bp38959547 to 38959618) |
| **H2347 A3A KO 11** | 50% | insertion of a T after H11 causing a frame shift and early stop codon in exon 3. there is also an insertion of a cg Y35 |
| | 50% | small deletion over guide 4 leading to premature stop codon in exon 4 |
| **H2347 A3A KO 12** | 50% | 20 bp deletion in guide 4 leading to frame shift and early stop codon in exon 4 |
| | 50% | 71 bp deletion leading to premature stop codon at cut site (aa12-36 exon 3; bp38959547 to 38959617) |
| **PC9 A3B KO 3** | 100% | Deletion: 1932 bp from 38984025 to 38985956. From guide 5 to guide 6. causes early stop codons in exon 3, 4, 5, 6 and 7 |
| **PC9 A3B KO 4** | 60% | Deletion: 2023 bp from 38983940 to 38985962. Early stop codons in 3, 4, 5, 6, and 7 |
| | 40% | Deletion: 1932 bp from 38984025 to 38985956. From guide 5 to guide 6. causes early stop codons in exon 3, 4, 5, 6 and 7 |
| **PC9 A3B KO 5** | 100% | Deletion: 1932 bp from 38984025 to 38985956. From guide 5 to guide 6. causes early stop codons in exon 3, 4, 5, 6 and 7 |
| **PC9 A3B KO 6** | 100% | Deletion: 1932 bp from 38984025 to 38985956. From guide 5 to guide 6. causes early stop codons in exon 3, 4, 5, 6 and 7 |
| **PC9 A3B KO 7** | 22.22% | Deletion: 2023 bp from 38983940 to 38985962. Early stop codons in 3, 4, 5, 6, and 7 |
| | 77.78% | Deletion: 1932 bp from 38984025 to 38985956. From guide 5 to guide 6. causes early stop codons in exon 3, 4, 5, 6 and 7 |
| **PC9 A3B KO 8** | 100% | Deletion: 1932 bp from 38984025 to 38985956. From guide 5 to guide 6. causes early stop codons in exon 3, 4, 5, 6 and 7 |
| **PC9 A3B KO 9** | 100% | Deletion: 1932 bp from 38984025 to 38985956. From guide 5 to guide 6. causes early stop codons in exon 3, 4, 5, 6 and 7 |
| **PC9 A3B KO 10** | 100% | Deletion: 1932 bp from 38984025 to 38985956. From guide 5 to guide 6. causes early stop codons in exon 3, 4, 5, 6 and 7 |
| **H1650 A3B KO 1** | 100% | deletion of first T in guide 6 leading to early stop codon in L 116 |
| **H1650 A3B KO 2** | 100% | deletion of g6 (TCCTGTCTGAGCACCC) causing frame shift and early stop codon right after in H111 in exon 3 |
| **H1650 A3B KO 3** | 100% | 10k bp deletion of the entirety of exon 2 and half of exon three until F107 |
| **H1650 A3B KO 4** | 33.33% | deletion of g6 (TCCTGTCTGAGCACCC) causing frame shift and early stop codon right after in H111 in exon 3 |
| | 66.67% | 10k bp deletion of the entirety of exon 2 and half of exon three until F107 |

| | | |
|---|---|---|
| **H1650 A3B KO 6** | 75% | deletion of g6 (TCCTGTCTGAGCACCC) causing frame shift and early stop codon right after in H111 in exon 3 |
| | 25% | deletion of first T in guide 6 leading to early stop codon in L116 |
| **H1650 A3B KO 10** | 75% | deletion of g6 (TCCTGTCTGAGCACCC) causing frame shift and early stop codon right after in H111 in exon 3 |
| | 25% | deletion of first T in guide 6 leading to early stop codon in L116 |
| **H2347 A3B KO 1** | 100% | Guide 6 deletion (TCCTGTCTGAGCACCC) causes early stop codon at aa110 |
| **H2347 A3B KO 3** | 100% | insertion of a T in front of F107 causing stop codon in aa110 |
| **H2347 A3B KO 4** | 100% | Deletion: 2023 bp from 38983940 to 38985962. Early stop codons in 3, 4, 5, 6, and 7 |
| **H2347 A3B KO 5** | 20% | Deletion: 2023 bp from 38983940 to 38985962. Early stop codons in 3, 4, 5, 6, and 7 |
| | 80% | Deletion: 1932bp from 38984025 to 38985956. Leads to the full deletion of exon 2, and premature stop codons in 3, 4, 5, 6 and 7 |
| **H2347 A3B KO 6** | 50% | deletion of g6 (TCCTGTCTGAGCACCC) causing frame shift and early stop codon right after in H111 in exon 3 |
| | 50% | (GTGTGGCGAAGCTGGCCGAATTCCTGTCTGA) del of g6 and a small upstream sequence causes early stop codon aa106 |

**Supplementary Table 1. APOBEC3 Targeted Sequence results.** Due to multiple chromosome copies, APOBEC3 knock-outs were confirmed by TOPO-cloning and Sanger sequencing of PCR products targeting the APOBEC3A and APOBEC3B loci.

| Plasmid Name | Lab ID/Addgene # |
|---|---|
| pU6-sgA3A.g1-Cas9-T2A-mCherry | pKC648 |
| pU6-sgA3A.g2-Cas9-T2A-mCherry | pKC649 |
| pU6-sgA3B.g1-Cas9-T2A-mCherry | pKC539 |
| pU6-sgA3B.g2-Cas9-T2A-mCherry | pKC540 |
| pU6-sgA3C.g1_CBh-Cas9-T2A-mCh | pJS1108 |
| pU6-sgA3C.g2_CBh-Cas9-T2A-mCh | pJS1109 |
| pU6-sgA3F.g1_CBh-Cas9-T2A-mCh | pJS1112 |
| pU6-sgA3F.g2_CBh-Cas9-T2A-mCh | pJS1113 |
| pU6-sgA3H.g1_CBh-Cas9-T2A-mCh | pJS1114 |
| pU6-sgA3H.g2_CBh-Cas9-T2A-mCh | pJS1115 |
| HP138-A3F-GFP | pAD727 |

**Supplementary Table 2. Plasmids used to target APOBEC loci.** Plasmid names and IDs used for generated APOBEC3 knockouts, knockdowns and APOBEC3F overexpression

| Primer Name | Primer Sequence | Lab Identifier |
|---|---|---|
| A3A F | ATGCTCGGTGTGGTAGGAGT | JM669 |
| A3A R | CCACAAGTACAATCCGGAACCT | JM670 |
| A3B F1 | TGTATTTCAAGCCTCAGTACCACG | JM679 |
| A3B R1 | CATAGTCCATGATCGTCACGC | JM680 |
| A3B F2 | CCTCAGAATTCAGTTCTACCATGA | JM663 |
| A3B R2 | TCGATTGAAGTTTGGGTGGTA | JM664 |
| A3B R3 | GAATCTCCTTTAGCGTGCGG | JM636 |

**Supplementary Table 3. Sequencing primers.** Primer names, sequences, and identifiers used for targeted sequencing.

| Gene | Forward Primer Sequence | Reverse Primer Sequence |
|---|---|---|
| *APOBEC3A* | gagaagggacaagcacatgg | tggatccatcaagtgtctgg |
| *APOBEC3B* | gaccctttggtccttcgac | gcacagccccaggagaag |
| *APOBEC3C* | agcgcttcagaaaagagtgg | aagtttcgttccgatcgttg |
| *APOBEC3D* | acccaaacgtcagtcgaatc | cacatttctgcgtggttctc |
| *APOBEC3F* | ccgtttggacgcaaagat | ccaggtgatctggaaacactt |
| *APOBEC3G* | ccgaggacccgaaggttac | tccaacagtgctgaaattcg |
| *APOBEC3H* | agctgtggccagaagcac | cggaatgtttcggctgtt |
| *HPRT1* | tgaccttgatttattttgcatacc | cgagcaagacgttcagtcct |
| *ACTIN* | atctggcaccacaccttctac | cagccaggtccagacgcagg |

**Supplementary Table 4. Primers for RT-qPCR.** Primer names, sequences, and identifiers used for RT-qPCR.
